# Oysters from five continents harbour a vast array of previously unknown RNA viruses, including many putative viruses of vertebrates

**DOI:** 10.64898/2026.09.29.755536

**Authors:** Kevin Xu Zhong, Amy M. Chan, Curtis A. Suttle

## Abstract

Oysters are vital to global aquaculture and marine ecosystems, yet, their associated RNA viruses remain largely unexplored. Here, we analysed ∼1,000 publicly available metatranscriptomic datasets from 13 oyster species across 14 countries and 5 continents to characterize the oyster RNA virome (ORV). We identified 1,515 distinct RNA viruses putatively representing 17 viral orders and 766 species-level viral operational taxonomic units (vOTUs), with ∼62% being previously undocumented putative species. These viruses not only populate extant branches of RNA virus evolution, but also represent previously unrecognized families and genera, and include viruses that putatively infect bacteria, fungi, protists, plants, invertebrates, and vertebrates. Based on sequence similarity, we found many calicivirus-like, orthoreovirus-like, astrovirus-like, and picornavirus-like viruses that do not infect invertebrates or microbes; thus, oysters accumulate diverse viruses from the surrounding water, and can potentially be interrogated as reporters of viruses circulating in the environment.

## Introduction

Oysters are ecologically and economically keystone species, and a pillar of global aquaculture^1–3^. An oyster can filter >100 L of seawater per day^4,5^, affecting nutrient cycling, habitat structure, and water quality^3^. Oysters also harbour bacteria, fungi, protists, and viruses^3,6–9^, as well as metazoans (e.g., worms, sponges, cestodes, trematodes, gastropods, and copepods)^10–14^; some of these are commensal, while others are consumed^15–18^. Beyond their ecosystem services, oysters support a global multibillion-dollar aquaculture industry^2,19^.

Despite the importance of oysters, the viruses associated with them are largely enigmatic, particularly for RNA viruses, which pose risks for oyster and public health. Human pathogens such as norovirus, Hepatitis A virus (HAV), and Hepatitis E virus (HEV), along with rotavirus, astrovirus, sapovirus, and aichivirus can bioaccumulate in oysters, and potentially cause diseases if raw oysters are consumed^20–36^. In addition, viruses can threaten oyster health, and have been associated with disease and mass-mortality events^37,38^. Thus, there is a strong need to define the oyster virome from both public and ecosystem-health perspectives. Despite their ecological, environmental and economic importance, research on RNA viruses associated with oysters is limited to a few geographic regions and focused primarily on Pacific oysters (*Magallana gigas*, formerly *Crassostrea gigas*) and Hongkong oysters (*Magallana hongkongensis*, formerly *Crassostrea hongkongensis*). These studies have identified ∼200 different RNA viruses^39–45^.

In recent years, far more viruses have been discovered through environmental sequencing (viromics) than by characterizing viral isolates, leading to a consensus statement that was adopted by the *International Committee on the Taxonomy of Viruses* (ICTV) to allow viral taxa to be established from sequence data alone^46^, as well as proposed standards for describing an uncultivated virus genome^47^. For RNA viruses, the marker of choice for identifying and classifying RNA viruses in environmental sequence data is RNA-dependent RNA-polymerase (RdRp)^42,48–50^.

Here, we mined data from nearly 1,000 publicly available oyster metatranscriptomes, representing 13 oyster species from 14 countries and five continents, and assembled contiguous sequences (contigs) in which RdRp sequences were identified and classified. Within this comprehensive oyster RNA virome (ORV), we identified 1,515 distinct RNA viruses putatively representing 17 viral orders and 766 species-level viral operational taxonomic units (vOTUs). Of these, ∼62% are previously undocumented putative species; thus, expanding the known RNA virus diversity in oysters by ∼4-fold. Below, we explore some of the remarkable findings revealed in this comprehensive catalogue of oyster-associated RNA virus sequences.

## Results

### Oyster-associated RNA viruses (ORV) catalog

We investigated 938 metatranscriptomes from 13 different oyster species across 14 countries and five continents (Supplementary Fig. 1; Extended Data Table 1), including ∼96 metatranscriptomes of wild (healthy) and farmed (diseased and dying) adult Pacific oysters sampled from British Columbia (BC), Canada^38^. Of these, 774 metatranscriptomes from at least ten oyster species have not been described for RNA viruses in the current RNA viromic databases, including Serratus^51^, RVMT^52^, IMG/VR4^53^, Tara Ocean^54^, and LucaProt^55^. Using RdRp sequences as a marker for RNA viruses, we identified 1,515 non-redundant contigs (>1 kb). These contigs varied in size from 1,000 to 41,709 bp (median = 1,796 bp), averaging 2,669 ± 2,448 bp in length (Fig. 1a). We established the oyster-associated RNA viruses (ORV) catalog by applying a threshold of >90% average nucleotide identity (ANI) across >75% alignment fractions (AF); these non-redundant RdRp-containing contigs were further classified into 766 species-level RNA virus populations (vOTUs).

**Fig. 1.**
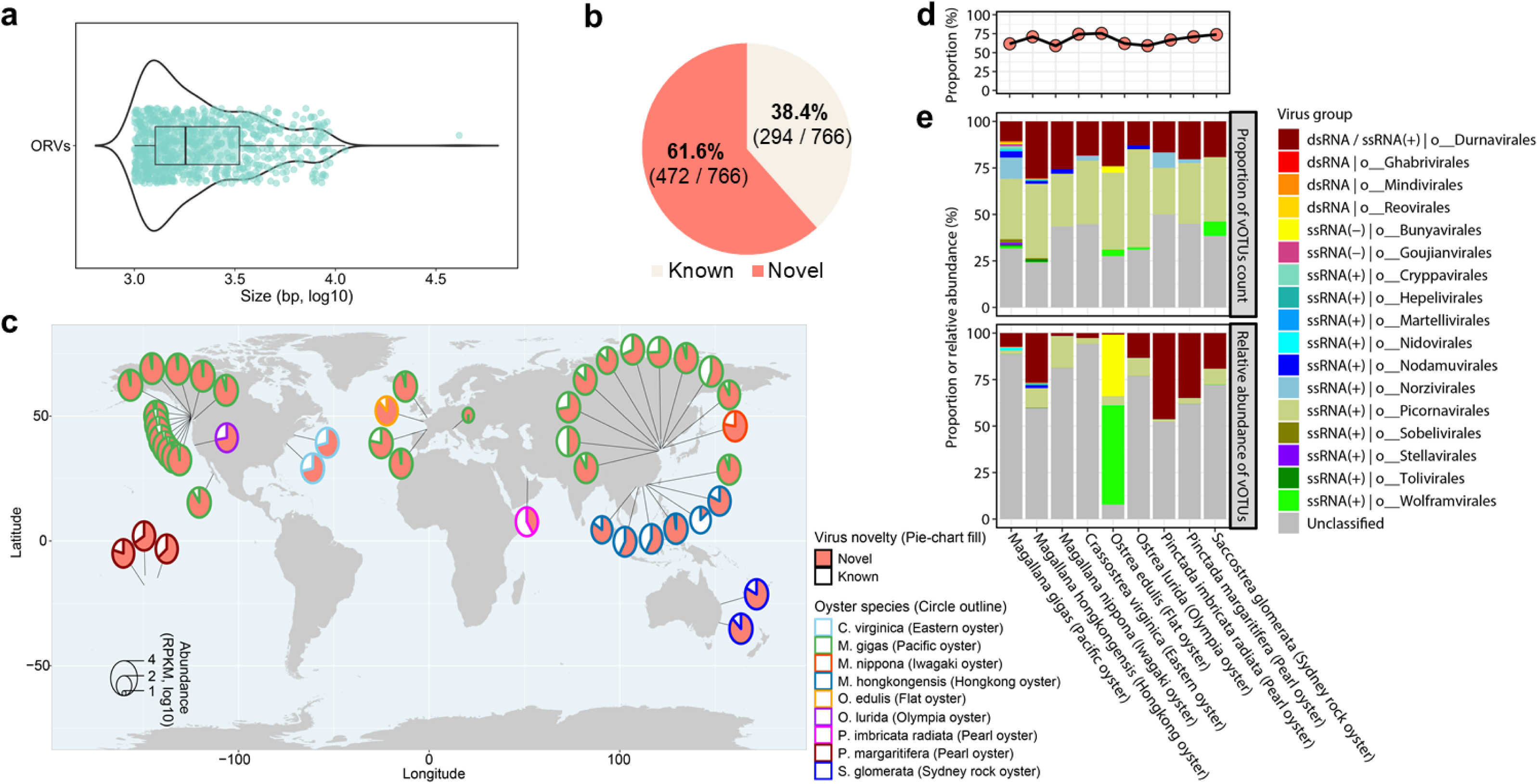
Detection of RNA viruses in global oyster populations. **a**, Genome size distribution of RdRp-containing contigs from 766 species-level (>90% ANI across >75% AF) RNA virus populations (vOTUs) identified in global oysters (Extended Data Tables 1 and 2). These oyster-associated RNA viruses (ORV) vOTUs are shown in light teal circles in the box plot. The box represents the interquartile range (IQR), with the line inside indicating the median value. The whiskers, extending from the left and right of the box, show the range within 1.5 times the IQR from the first and third quartiles. On the top of the box plot, a smoothed density plot of genome sizes is displayed, with the plot width reflecting the frequency of observations within each range of values. **b**, Novelty of RNA viruses in the ORV catalog. The vOTUs with high species-level genomic similarity (>90% ANI across >75% AF) to RNA virus sequences in NCBI RefSeq (release 216), Serratus^51^, Tara Ocean^54^, IMG/VR4^53^, RVMT^52^, and LucaProt^55^ databases are classified as “known”, while those with lower similarities are categorized as “novel”. **c**, Distribution of known and novel vOTUs across various oyster species and locations. Each circle represents a sampling location, with the outline color indicating the oyster species and circle size proportional to the RPKM-normalized abundance of RNA virus vOTUs detected at that location. Viral ‘abundance’ was determined by mapping metatranscriptomic reads to ORV vOTU genomes at >98% average nucleotide identity (ANI), followed by normalization to reads per kilobase per million (RPKM). The circle’s pie-chart fill indicates virus novelty (known or novel). Locations where no RNA viruses were detected are excluded from the visualization (n = 20; see details in Extended Data Table 3). **d**, Proportion of the number of novel vOTUs across oyster species. **e**, Order-level taxonomy distribution of vOTUs found across oyster species. The upper panel shows the proportion of the number of vOTUs in each oyster species, while the lower panel displays the relative abundance of vOTUs. In the panel **c**, **d** and **e**, oyster species without RNA viruses detected are excluded from the visualization (n = 4; see details in Supplementary Fig. 1b and Extended Data Table 3).

Of these 766 ORV vOTUs, 13 (1.7%), 27 (3.5%), 138 (18%), 340 (44.4%), and 248 (32.4%) were classified as complete, high-quality (>90% complete), medium-quality (50–90% complete), low-quality (<50% complete), and quality-not-determined genomes, respectively (Extended Data Table 2). These sequences were assembled from nine out of 13 oyster species, across eight out of 14 countries, and across four out of five continents (Supplementary Fig. 1b-c). Most ORV vOTUs (472, 61.6%) were derived from the metatranscriptomic datasets from the Hongkong oyster (*M. hongkongensis*; Supplementary Fig. 1b), while 28.5% (218) were from the Pacific oyster (*M. gigas*) and 6% (46) from the Olympia oyster (*Ostrea lurida*) datasets. A relatively small fraction (<2%) was obtained from datasets for the Eastern oyster (*Crassostrea virginica*), Iwagaki oyster (*Magallana nippona*, formerly *Crassostrea nippona*), Flat oyster (*O. edulis*), Sydney rock oyster (*Saccostrea glomerata*), and Pearl oysters in the species *Pinctada margaritifera* and *P. imbricata radiata*. Non vOTUs were assembled from Portuguese oysters (a.k.a., Fujian oysters: *Magallana angulata*, formerly *Crassostrea angulata*), Mangrove oysters (*Crassostrea tulipa*, formerly *Crassostrea gasar*), Australian black-lip oyster (*Saccostrea sp.* non-mordax lineage J), and Pearl oyster in the species *Pinctada fucata*.

### Discovery of novel RNA viruses in oysters

Of the 766 ORV vOTUs identified, 472 (61.6%) did not exhibit species-level nucleotide sequence similarity (>90% ANI across >75% AF) with sequences from RNA virome databases (Fig. 1b-d; Supplementary Fig. 2), highlighting the novelty for a substantial proportion of the RNA viruses in oysters. Specifically, only 2.1% (16), 14.5% (111), 9.8% (75), 30.9% (237), 22.1% (169), and 1% (8) of ORV vOTUs showed species-level nucleotide sequence similarity (>90% ANI across >75% AF) to RNA viruses from the NCBI RefSeq (release 216), IMG/VR4^53^, RVMT^52^, Serratus^51^, LucaProt^55^, and Tara Ocean^54^ databases, respectively (Supplementary Fig. 2a). Remarkably, only one ORV vOTU had species-level relatives across all six databases that include RNA virus sequences from a variety of hosts and ecosystems. Collectively, these findings show that oysters harbour a vast array of previously undescribed RNA viruses.

We assigned taxonomy to ORV vOTUs by leveraging the power of several virus classification pipelines, and further refined and curated the taxonomy using phylogenetic analysis of RdRp-containing polyprotein sequences (see Methods for details) (Figs. 1e and 2; Supplementary Figs. 3-12; Extended Data Table 2). All 766 ORV vOTUs were assigned to RNA virus in the realm *Riboviria*. Of these, ∼74.8% (573 vOTUs) could be classified at the kingdom level, whereas the remaining 193 vOTUs, comprising zhaoviruses, yanviruses, and weiviruses, were unclassified at this rank. The 573 kingdom-assigned vOTUs belonged to *Orthornavirae* and were further assigned to five phyla, 15 classes, and 17 orders (Figs. 1e and 2; Supplementary Figs. 4-12).

**Fig. 2.**
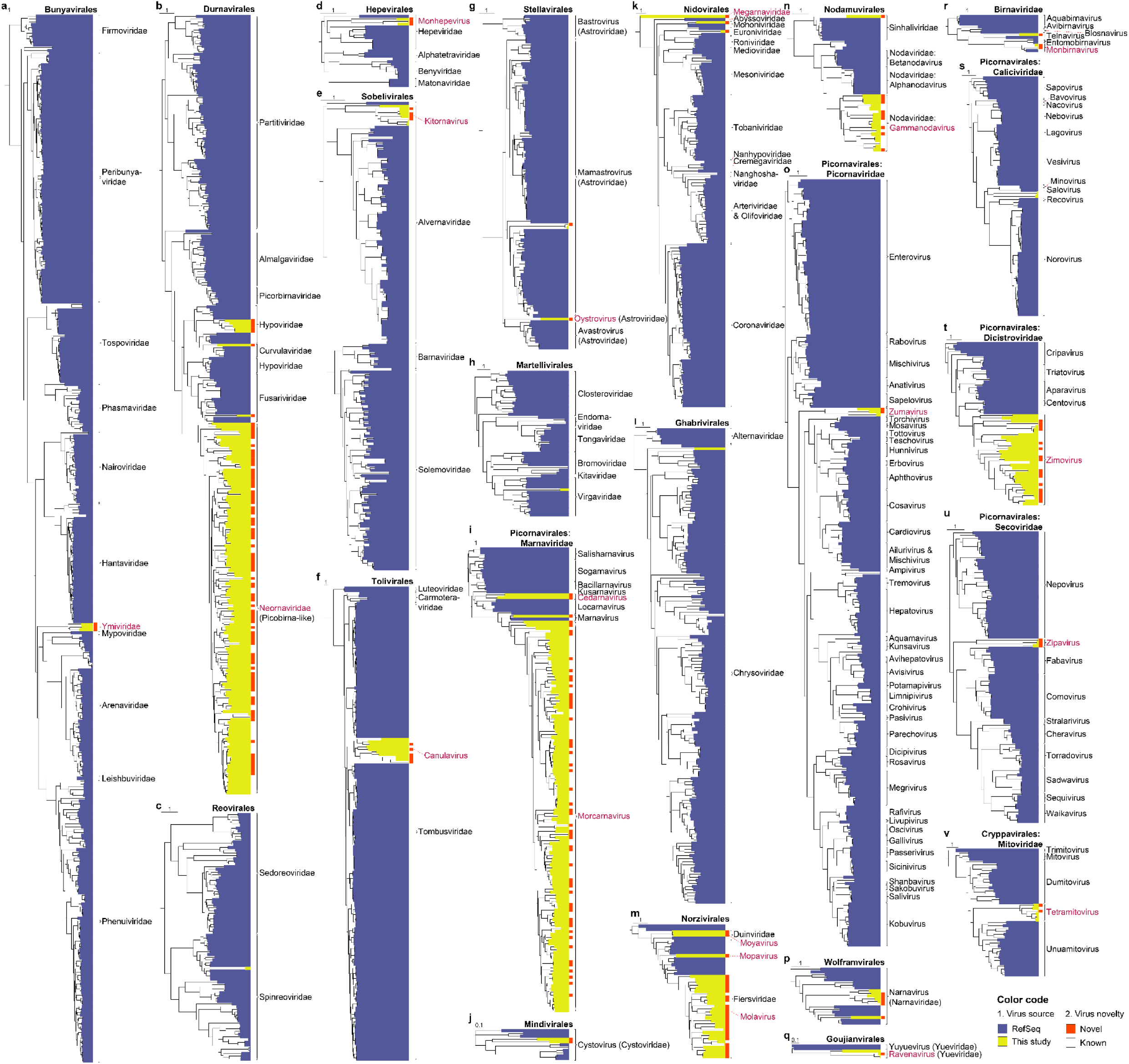
Phylogeny of RNA viruses detected in oysters. **a-v**, Phylogeny of RNA viruses across different viral orders (**a-q**, **s-v**) and viral families that are unclassified at the order level (**r**). Unrooted maximum-likelihood (ML) phylogenetic tree, based on analysis of the inferred amino-acid sequences of the polyprotein containing RNA-dependent RNA polymerase (RdRp), illustrates the evolutionary histories of oyster-associated RNA viruses (ORV) vOTUs (yellow) and other RNA viruses found in the NCBI RefSeq (release 216; blue). Tree leaves are colored to indicate whether the virus genome sequences share similarity (>90% ANI across >75% AF) with known sequences (white) or represent novel sequences (red) not found in NCBI RefSeq (release 216) or other RNA virome databases, including Serratus^51^, Tara Ocean^54^, IMG/VR4^53^, RVMT^52^, and LucaProt^55^. The tree scale bar represents the relative rate of evolution along the branches of the ML tree. Tree leaves are annotated based on the established taxonomy, clustering pattern, and bootstrap support values (see details in Supplementary Figs. 4-12); text highlighted in blue indicates proposed families or genera names for each viral group.

Based on phylogenetic analysis, over half (434/766; 56.7%) of the ORV vOTUs are sufficiently divergent from RNA viruses in the NCBI RefSeq (release 216) that putatively, they represent three new families and 16 novel genera (Fig. 2; Supplementary Figs. 4-12; see Supplementary Results for details). These viruses represent new putative families or genera (e.g., *Ymiviridae*, *Neornaviridae*, *Oystrovirus*, *Cedamavirus*, *Zipavirus*, *Zumavirus*, *Monhepevirus*, *Canulavirus*, *Tetramitovirus*, *Mopavirus*, and *Moyavirus*) that occupy positions between established phylogenetic branches, or are on deep branches within established phylogenetic groups (e.g., the proposed family *Megarnaviridae*, and genera *Ravenavirus*, *Kitornavirus*, *Zinmovirus*, *Morcamavirus*, *Gammanodavirus*, *Monbirnavirus*, and *Molavirus*).

### Diverse probable hosts for oyster-associated RNA viruses

Based on phylogenetic relationships to viruses with known hosts, we predict the ORV vOTUs represent viruses that infect an array of hosts, including bacteria, fungi, protists, algae, higher plants, invertebrates, and vertebrates (Fig. 3). Based on ORV vOTUs, bacteria, fungi, and plants/protists/algae are predicted to be likely hosts for 4.3% (33), 2.7% (21), and 20.5% (157), respectively, of the predicted viruses; whereas, invertebrates and vertebrates are predicted hosts for 8.1% (62) and 4.3% (33) of the viruses, respectively. Putative hosts could not be determined for the remaining ∼60% of the predicted viruses, including members of *Monhepevirus* (*Hepeviridae*; order *Hepelivirales*), *Neornaviridae* (order *Durnavirales*), *Ymiviridae* (order *Bunyavirales*), unclassified *Picornavirales*, and other deeply divergent RNA viruses within *Orthornavirae* (Fig. 3; Extended Data Table 2; Supplementary Figs. 4a, 6a and 10a). This indeterminacy stems either from phylogenetic divergence, which obscures probable host predictions, or from the unresolved host relationships of their closely related RNA viruses.

**Fig. 3.**
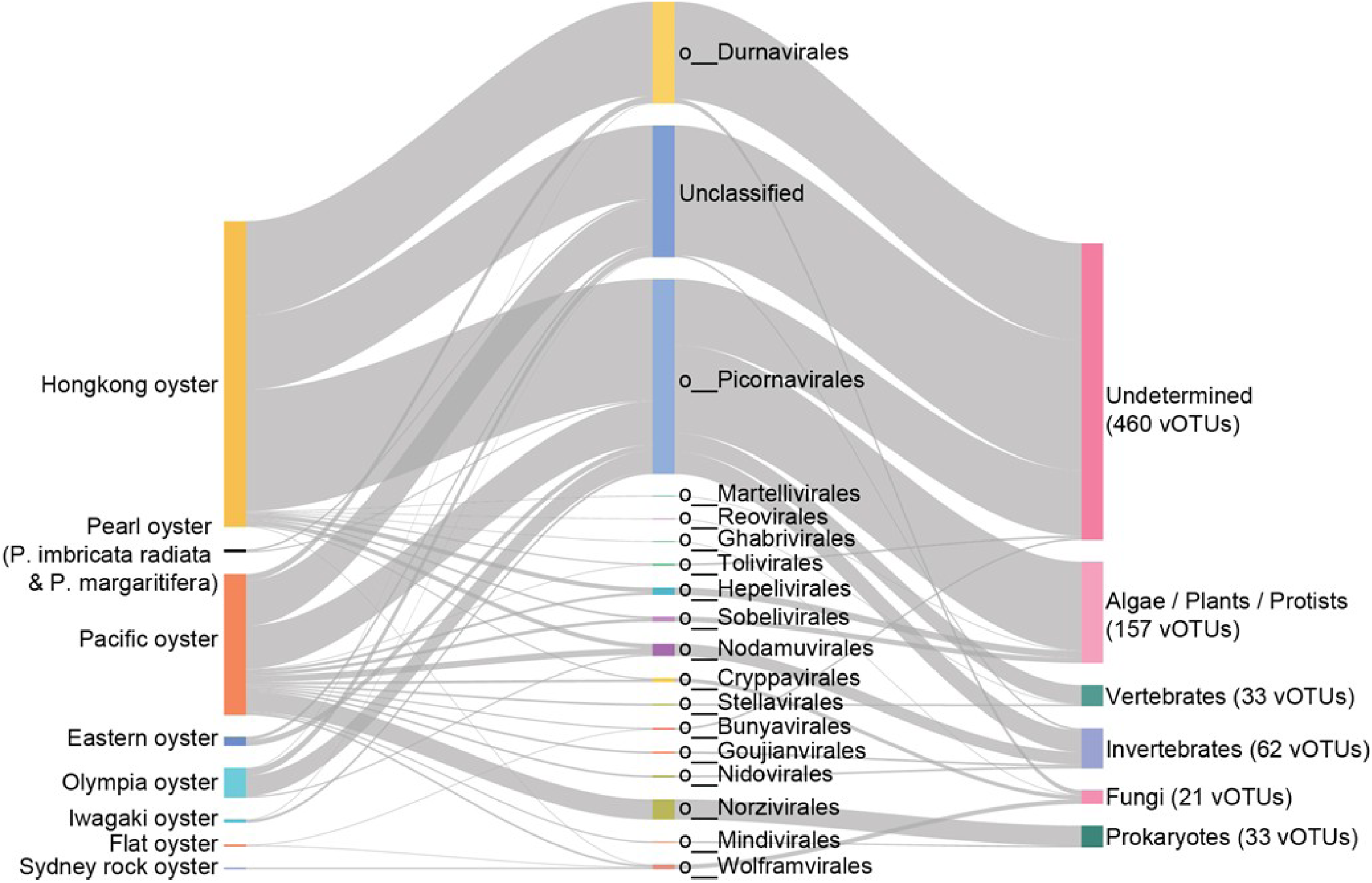
Putative hosts of oyster-associated RNA viruses (ORV) vOTUs. The plot illustrates the relationships between oysters (left column), virus orders of the ORV vOTUs (middle column), and their putative hosts (right column), interconnected by grey lines. Numbers in parentheses indicate the number of ORV vOTUs associated with each host group.

### Invertebrate-infecting RNA viruses in oysters

Invertebrate-infecting RNA viruses are diverse and widespread across oyster species, spanning positive-sense single-stranded RNA [(+)ssRNA], negative-sense single-stranded RNA [(-)ssRNA], and double-stranded RNA (dsRNA) viruses (Fig. 3). They included members of *Birnaviridae*, *Nodaviridae* (order *Nodamuvirales*), *Sinhaliviridae* (order *Nodamuvirales*), *Dicistroviridae* (order *Picornavirales*), *Yueviridae* (order *Goujianvirales*), and several families in the *Nidovirales* (*Megarnaviridae*, *Mononiviridae*, and *Euroniviridae*) (Supplementary Figs. 4b, 5a, 7a, 9b, and 11a). Viruses in these taxa are known to infect invertebrates, and thus those found in this study likely infect oysters or other oyster-associated invertebrates, such as, worms, copepods, nematodes, trematodes, gastropods and sponges^10–14^. In addition, small invertebrates including zooplankton consumed as food^15–18^, as well as free viral particles^56^ from the surrounding seawater are other potential sources of these viruses.

Several RNA viruses were identified that likely infect oysters. For example, Pacific Oyster Nidovirus 1 (PONV1; ORV316), a member of the tentative genus *Alphamegarnavirus* within the tentative family *Megarnaviridae* (order *Nidovirales*), was widespread in Pacific oysters globally and has been associated with mass mortality in farmed oysters^38^ (Fig. 4; Supplementary Figs. 5a and 13). Three dsRNA *Birnaviridae*-vOTUs (ORV574–576) were also detected. ORV574, detected in Hongkong oysters from the Shenzhen region in China, clustered with bivalve-infecting Tellina virus 1 (TV-1)^57^ in the genus *Telnavirus* (Fig. 4; Supplementary Fig. 11a). In contrast, ORV575 and ORV576 grouped with *Pinctada birnavirus* TM-2022 (PiBV)^58–61^, but in a distinct clade tentatively designated *Monbirnavirus*; these were detected in a few Pacific oyster samples from BC, Canada (Fig. 4; Supplementary Figs. 11a and 13).

**Fig. 4.**
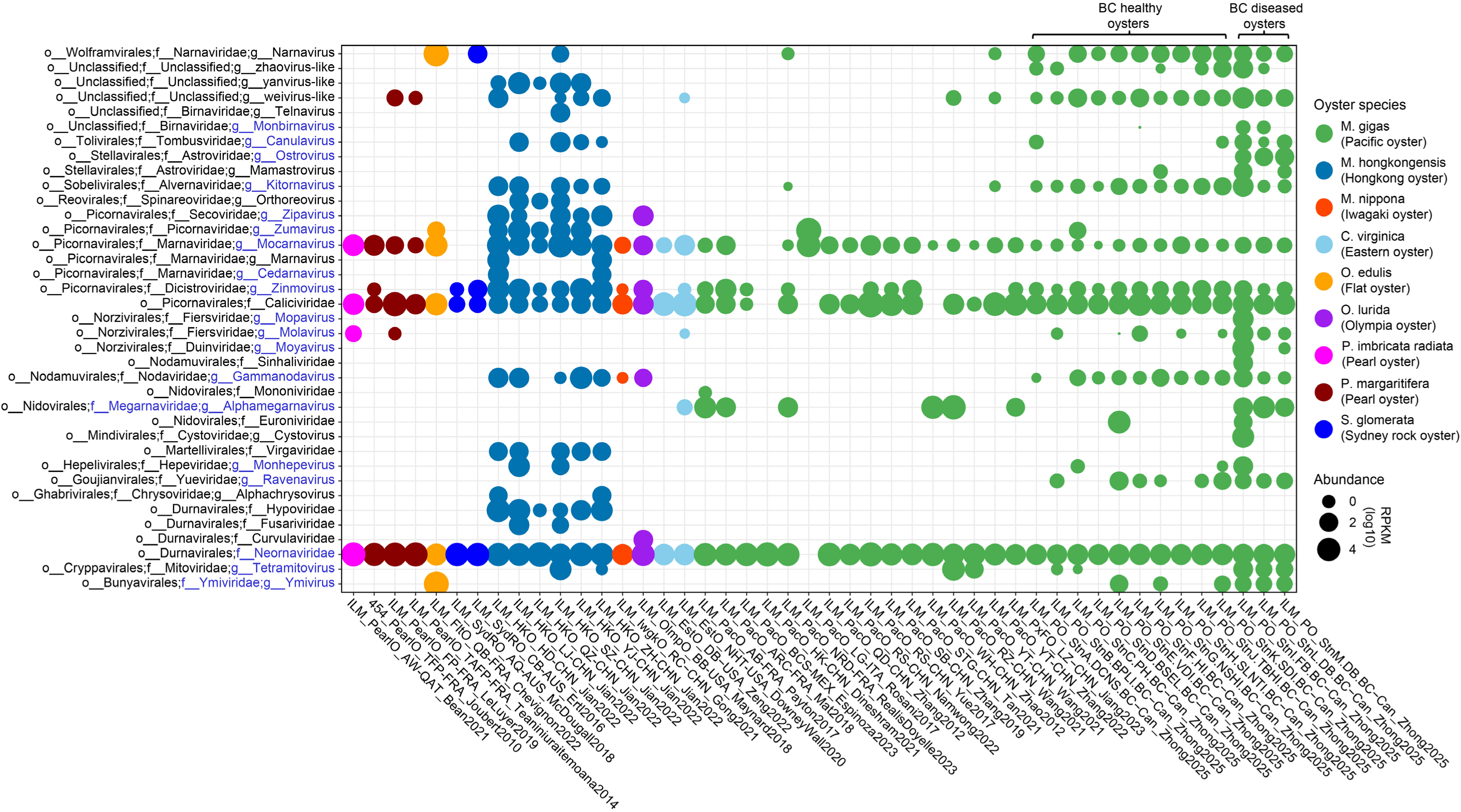
Prevalence of oyster-associated RNA viruses assigned to newly proposed families and genera in oysters across locations. Each circle represents the detection of viruses, with the circle size indicating the RPKM-normalized abundance of these viruses in oysters from each location (n = 46). The RPKM-normalized abundance was determined by mapping metatranscriptomic reads to the newly proposed viral families or genera with an average nucleotide similarity (ANI) >98% and then normalizing the mapped reads to RPKM (reads per kilobase per million). It provides a semi-quantitative assessment of the examined viruses present in oysters across various locations, revealing the relative levels of viral loads in the host. Locations where no RNA viruses were detected are excluded from the visualization (n = 20; see details in Extended Data Table 3).

Moreover, RNA viruses related to those infecting oyster-associated invertebrates, including worms (ORV314; family *Mononiviridae*, order *Nidovirales*) and arthropods (ORV315; family *Euroniviridae*, order *Nidovirales*), were identified, suggesting acquisition from associated fauna or from invertebrate-derived viral particles from the surrounding seawater (see Supplementary Results). Additional invertebrate-infecting RNA viruses were also present, though it is unclear whether their hosts are oysters, oyster-associated invertebrates, or other environmental invertebrates. These include the tentative sinaiviruses (*Sinhaliviridae*) and gammanodaviruses (*Nodaviridae*) within the *Nodamuvirales*, zinmoviruses (*Dicistroviridae*; *Picornavirales*), and ravenaviruses (*Yueviridae*; *Goujianvirales*) (Supplementary Figs. 4b, 7a, and 9b). Zinmoviruses and gammanodaviruses were widespread across oyster species and worldwide, suggesting a long-term ecological and/or evolutionary association with oysters (Fig. 4). By contrast, sinaiviruses and ravenaviruses were geographically restricted and host-specific. For example, they were detected exclusively in Pacific oysters from BC, Canada, likely reflecting localized oyster-associated fauna or environmental conditions (see details in Supplementary Results).

### Microbe-associated and food-derived RNA viruses in oysters

We also detected RNA viruses likely infecting oyster-associated microbes (Fig. 3). These included bacteriophages in the families *Fiersviridae* and *Duinviridae* (order *Norzivirales*) and *Cystoviridae* (order *Mindivirales*), as well as fungal viruses in the families *Mitoviridae* (order *Cryppavirales*), *Narnaviridae* (order *Wolframvirales*), *Crysoviridae* (order *Ghabrivirales*), and several *Durnavirales* families (*Hypoviridae*, *Fusaviridae*, and *Curvulaviridae*) (Supplementary Figs. 6a, 11, and 12). Many of these bacteriophages and fungal viruses may be derived from free virus particles in seawater or associated with other organisms (e.g., phytoplankton, microzooplankton, protists) within oysters. Most RNA bacteriophages, including cystoviruses (order *Mindivirales*) and tentative mopaviruses and moyaviruses (order *Norzivirales*) (e.g., *Cystovirus* in *Mindivirales*, and *Mopavirus* and *Moyavirus* in *Norzivirales*), were only detected in adult Pacific oysters in BC, occurring in both healthy and diseased individuals (Fig. 4; Supplementary Fig. 3). *Molavirus*, however, was also observed in Pearl and Eastern oysters. Fungi-infecting RNA viruses showed a similarly heterogeneous distribution: *Crysoviridae*, *Hypoviridae*, and *Fusaviridae* were found only in Hongkong oysters, whereas *Mitoviridae* and *Narnaviridae* were found in Pacific, Flat, and Sydney rock oysters.

The second most prevalent RNA viruses in oysters were likely associated with plant material including phytoplankton, and microzooplankton (Fig. 3); they accounted for >20% of ORV vOTUs and included viruses in the families *Marnaviridae* (order *Picornavirales*), *Secoviridae* (order *Picornavirales*), *Avelrnaviridae* (order *Sobelivirales*), *Tombusviridae* (order *Tolivirales*), and *Vigaviridae* (order *Martellivirales*) (Supplementary Figs. 5c, 7, 9a, and 10b).

### Vertebrate-infecting RNA viruses in oysters

A subset of the ORV vOTUs (33 of 766) were assigned to taxa with viruses known to infect vertebrates and were detected across multiple oyster species at various locations (Fig. 4). These include (+)ssRNA viruses in the families *Caliciviridae* (order *Picornavirales*), *Picornaviridae* (order *Picornavirales*), and *Astroviridae* (order *Stellavirales*), as well as dsRNA viruses in the family *Spinareoviridae* (order *Reovirales*). The presence of viruses in oysters that are only known to infect vertebrates implies that they are accumulated from seawater (Supplementary Figs. 5b, 6b, 8a, and 11c).

*Caliciviridae*-like vOTUs (ORV222 and ORV301) encode a putative calicivirus coat protein domain (InterPro entry: IPR029053) and form a sister clade (bootstrap value >50%) with known fish caliciviruses, Atlantic salmon calicivirus (genus *Salovirus*)^62^ and Fathead minnow caliciviruses (genus *Minovirus*)^63^ in phylogenies generated for both RdRp-containing polyprotein and RdRp palm sequence domains (Supplementary Figs. 6b and 15; Fig. 2). These oyster-associated caliciviruses were widespread, occurring in all nine-oyster species across almost all locations (43 out of 46) where RNA viruses were detected, as well as in all farmed (diseased) and wild (healthy) adult Pacific oysters in BC (Fig. 4; Supplementary Fig. 13). Notably, vOTU ORV222, comprises nine distinct calicivirus contigs assembled from geographically widespread oyster metatranscriptomes, indicating substantial within-species viral diversity across locations and oyster species. In contrast, vOTU ORV301 is represented by a single calicivirus contig, showing a more restricted distribution, largely confined to Flat oysters from China (Fig. 5).

**Fig. 5.**
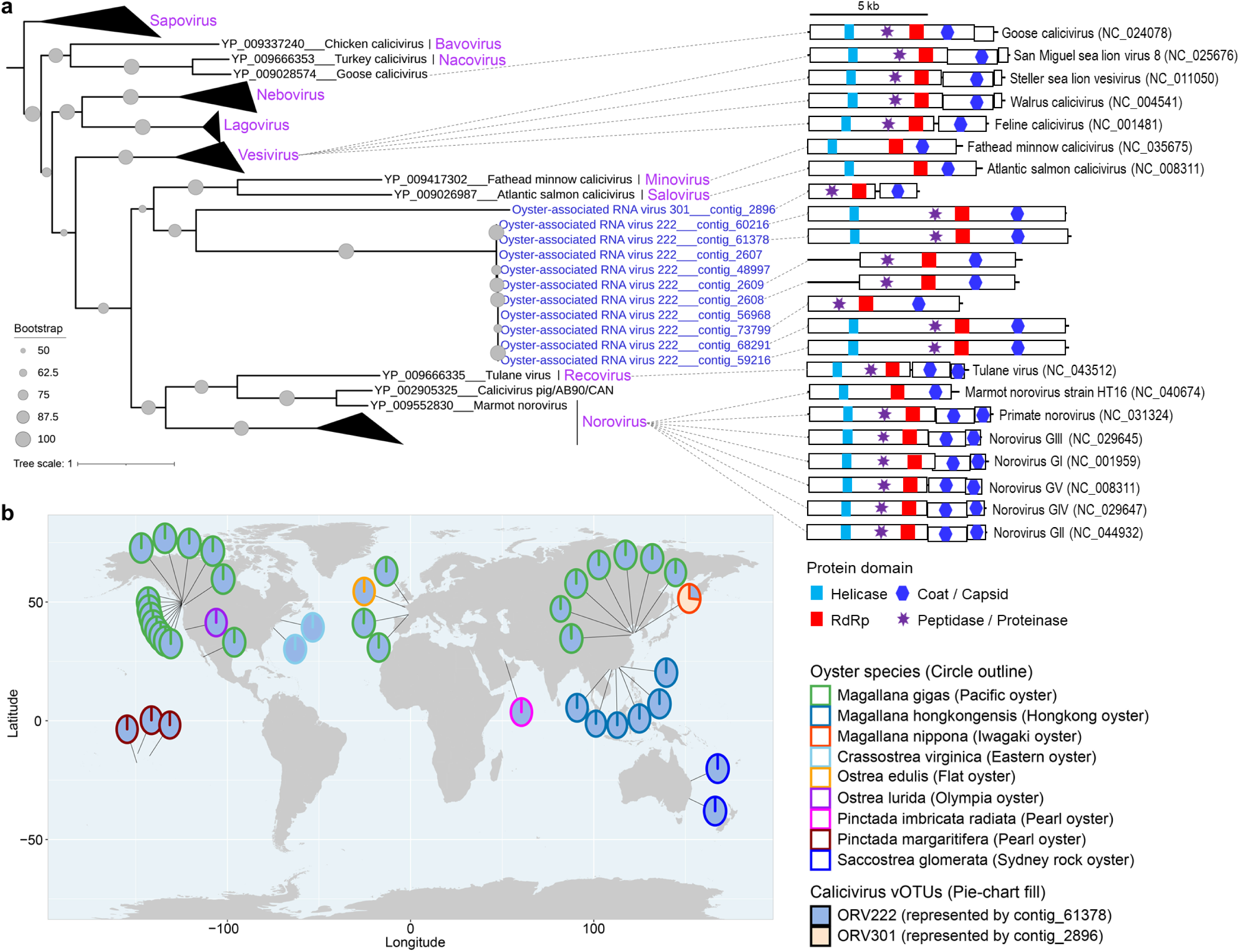
Detection of caliciviruses in oysters across various species and locations. **a**, Phylogenetic placement of oyster-associated caliciviruses. The maximum-likelihood (ML) phylogenetic tree, based on the inferred amino acid sequence of the polyprotein containing RNA-dependent RNA polymerase (RdRp), illustrates the evolutionary relationships between caliciviruses detected in oysters (blue text at tree tips) and those from the NCBI RefSeq (release 216; black text at tree tips). Node labels indicate bootstrap support values. The tree is grouped based on shared taxonomy at genus level, labeled in pink. Representative virus genome map with annotated protein domains is displayed adjacent to the tree leaf, connected by a grey dashed line. For the genome map of oyster-associated caliciviruses, only those with >4 kb are shown. The ML tree is re-rooted using members of the genus *Sapovirus*. **b**, Relative composition of calicivirus vOTUs among the total oyster-associated caliciviruses across various oyster species and locations. Each circle represents a location at which caliciviruses were detected, with its outline colored according to oyster species. Pie charts within circles show the relative composition of calicivirus vOTUs, calculated as the RPKM abundance of each vOTU divided by the summed RPKM abundance of all calicivirus vOTUs detected at that location. Circle size is fixed and does not represent viral abundance. RPKM values were calculated by mapping metatranscriptomic reads to calicivirus genome sequences at >98% average nucleotide identity (ANI) and normalizing mapped reads as reads per kilobase per million (RPKM). Locations where no caliciviruses were detected are excluded from the visualization (n = 23; see details in Fig. 4 and Extended Data Table 3).

Other vertebrate-infecting RNA viruses included *Orthoreovirus* (*Spinareoviridae*; *Reovirales*), *Mamastrovirus*, and the tentatively assigned *Oystrovirus* (*Astroviridae*; *Stellavirales*) and *Zumavirus* (*Picornaviridae*; *Picornavirales*) (Supplementary Figs. 5b, 8a, and 11c). These viruses tended to show more restricted geographic distributions, suggesting regional environmental sources or localized contamination. For example, mamastroviruses and oystroviruses were only found in Pacific oysters in BC, Canada, while orthoreoviruses were only detected in Hongkong oysters in China (Fig. 4; Supplementary Fig. 13). See details in Supplementary Results.

### RNA viruses with host undetermined

Finally, a large subset of ORV vOTUs (460 of 766) could not be assigned to a putative host (Fig. 3; Extended Data Table 2). These include tentatively assigned *Neornaviridae* (order *Durnavirales*), *Ymiviridae* (order *Bunyavirales*), and *Monhepevirus* (family *Hepeviridae*; order *Hepelivirales*) (Fig. 3; Supplementary Figs. 4a, 6a, and 10a). Notably, 137 vOTUs (17.9% of all ORVs) were *Picobirnaviridae*-like vOTUs placed within the proposed family *Neornaviridae* (Fig. 3; Supplementary Fig. 6a). These *Neornaviridae* vOTUs have *Picobirnaviridae*-like RdRp sequences that are phylogenetically branched between the vertebrate-infecting *Picobirnaviridae* and fungi-infecting viruses of the order *Durnavirales* (e.g., *Curvulaviridae*, *Hypoviridae*, and *Fusariviridae*), a pattern also noted previously^40,41,64^. Although their hosts are unknown, recent evidence suggests that such lineages may infect bacteria^64^. Noteworthy, these *Neornaviridae* (*Picobirnaviridae*-like viruses) were widespread, occurring in nearly all oyster species and locations, including all farmed and wild Pacific oysters in BC (Fig. 4; Supplementary Fig. 13).

Similarly, probable hosts of the *Ymiviridae* and *Monhepevirus* remain unresolved. Both groups occupy branching positions between viral lineages known to infect different host types, making host assignment ambiguous. Members of *Ymiviridae* and *Monhepevirus* were both present in farmed (diseased) and wild (healthy) Pacific oysters in BC. As well, *Ymiviridae* also occurred in Flat oysters from France, and *Monhepevirus* was also detected in Hongkong oysters across several regions in China (Fig. 4; Supplementary Fig. 13). See details in Supplementary Results.

## Discussion

Oysters hold significant ecological and economic value as key species in global aquaculture and marine ecosystems. Documenting the RNA viruses associated with oysters thus holds substantial promise for understanding broader ecological dynamics, pathogen transmission, and aquaculture sustainability. This study mined RNA virus diversity in publicly available metatranscriptomic data from 13 oyster species across 14 countries and five continents (Supplementary Fig. 1), and catalogued 1,515 different RNA viruses, organized into 766 species-level vOTUs. These viruses span 17 established orders, representing an approximately four-fold increase in the known RNA virus diversity of oysters (Figs. 1-3; Supplementary Figs. 4-12). Notably, over ∼62% (472/766) of these sequences belong to previously undescribed taxa, including putative new families and genera, and highlight a complex, rich, and largely unknown viral assemblage in oysters, including many viruses that likely infect plants and other animals. These findings imply that oysters accumulate a wide array of viruses from the surrounding seawater, and thus may serve as sentinels of coastal viral communities and as bioindicators of environmental contamination, with potential roles in environmental virus removal and contributions to overall ecosystem health. These observations and their implications are discussed in detail below.

The high diversity of viruses in oysters reflects the complex assemblage of metazoans (e.g., worms) and microbiota (e.g., prokaryotes, fungi, protists, viruses) that are resident in or consumed by oysters^6–18^, which includes potential viral hosts, as well as viruses that infect the oysters themselves. This diverse input pool yields a rich virome that includes 1,515 distinct RNA viruses, representing 766 putative viral species (vOTUs), with individual samples containing up to 46 distinct RNA viruses (RdRp-containing contigs >1 kb). The ORV comprises the richest viral diversity of any aquatic animal surveyed to date, and exceeds that documented in fishes (47-186 vOTUs)^65,66^, crustaceans (90 vOTUs)^67^, and in 58 aquatic invertebrate species (363 vOTUs)^41^. Although this number is lower than reported for terrestrial bats (∼4,000)^68,69^ and ticks (1,801)^70^, the ORV is one of the largest inventories of RNA viruses in animals, noting that quantitative comparisons across studies are constrained by differences in sampling scale, sequencing depth, and virus discovery criteria. Nonetheless, rarefaction analyses indicate that the diversity of RNA viruses in oysters remain under-sampled, and is even greater than presented here (Supplementary Fig. 14). Consistent with this, a large fraction of ORV vOTUs lack taxonomic classification (193 of 766) or host assignment (460 of 766), which highlights the extent of unexplored viral diversity harboured by oysters and suggests the need for targeted virus isolation and experimental studies to resolve viral taxonomy, host range, and ecological function. Additional complexity is introduced by the use of non-standard genetic codes that was inferred for ∼3.9% of ORV vOTUs (30 of 766; Extended Data Table 2), hinting at further uncharacterized aspects of RNA virus biology.

The high proportion of ORV vOTUs (∼62%) without close relatives in extant databases highlights how little of the marine RNA virosphere has been characterized in oysters. These novel vOTUs were spread across all 17 orders of RNA viruses identified in this study and were broadly distributed across locations, oyster species, development stages (e.g., larvae, juveniles, or adults), and health status (e.g., healthy, diseased, or under stress) (Figs. 1-2; Extended Data Tables 1-2; Supplementary Figs. 2-12). These data potentially establish three new viral families and 16 novel genera, demonstrating that oysters harbour a milieu of RNA viruses that previously were largely unknown. Much of this diversity likely stems from the wide range of particulate material ingested by oysters, resulting in the acquisition and accumulation of environmental viruses.

A notable component of the ORV is the relatively large number (62 vOTUs) of putative invertebrate-infecting viruses, such as those in the families *Birnaviridae* (*Monbirnavirus* and *Telnavirus*), *Nodaviridae* (*Sinaivirus* and *Gammanodavirus*), *Mononiviridae*, *Euroniviridae*, *Megarnaviridae* (*Alphamegarnavirus*), and *Dicistroviridae* (*Zinmovirus*). These likely include viruses that infect oysters, associated fauna (e.g., worms) and planktonic food (e.g., crustacean zooplankton). Thus, oysters sit at the nexus of microbial, planktonic, and metazoan interactions, and their viromes reflect this ecological integration.

Several of the viruses identified here have potential implications for oyster health. PONV1 (ORV316), in the proposed *Megarnaviridae*, has been linked to mass mortality in Pacific oysters in BC^38^; as well, two *Birnaviridae*-like vOTUs (ORV675-576 in the proposed genus *Monbirnavirus*) are closely related to PiBV, a virus associated with summer atrophy in Japanese Pearl oysters^58–61^ (Fig. 2; Supplementary Figs. 5a and 11a). Additional viruses, including widespread gammanodaviruses and zinmoviruses, and geographically restricted sinaiviruses and ravenaviruses, further illustrate the complexity of oyster-associated viral communities (Figs. 2 and 4; Supplementary Figs. 4b, 7a, 9b, 10a and 13). Collectively, these viruses highlight the diverse and dynamic roles oysters may play in marine viral transmission networks and the implications for ecosystem function, aquaculture sustainability, and invertebrate biodiversity.

Oysters also accumulate viruses related to those causing disease in humans and other vertebrates. Putative vertebrate-infecting RNA viruses were detected across multiple oyster species and locations, reflecting viral inputs from diverse environmental sources, including sewage and runoff, as well as vertebrates such as fish, and other coastal wildlife. These include viruses clustering with fish-associated caliciviruses (salovirus and minovirus), but not with human noroviruses, which are frequent contaminants of oysters^23–35,71–75^. These oyster-associated caliciviruses were widespread, occurring in nine oyster species across 43 of 66 sampling locations (Fig. 4; Extended Data Table 2), indicating efficient accumulation of environmental caliciviruses. Notably, ORV222 showed a near-global distribution across oyster species and regions, whereas ORV301 displayed a more restricted pattern, largely associated with Flat oysters detected in China, suggesting differences in environmental prevalence or region-specific sources (Fig. 5). Although these caliciviruses are not closely related to known human pathogens and their capacity to infect other vertebrates remains unknown, their pervasive presence indicates that oysters commonly accumulate animal-associated viruses from surrounding waters, consistent with reports of caliciviruses infecting a range of marine vertebrates, including fish^62,63^, sea lions^76–78^, and walruses^79^.

In contrast to caliciviruses, other vertebrate-associated RNA viruses, including those in the genera *Orthoreovirus* (*Spinareoviridae*, *Reovirales*), *Mamastrovirus* and *Oystrovirus* (*Astroviridae*, *Stellavirales*), and *Zumavirus* (*Picornaviridae*, *Picornavirales*), showed more restricted distributions across oyster species (e.g., Hongkong, Pacific, and/or Flat oysters) and locations (Fig. 4; Supplementary Figs. 5b, 8a and 11c). For example, mamastroviruses (ORV2-3) detected in Pacific oysters echo their presence in marine mammals such as pinnipeds, sea lions, and dolphins^80,81^. Similarly, oystrovirus (ORV1) and orthoreovirus (ORV572) cluster closely with avian viruses in the families *Astroviridae* and *Spinareoviridae*, respectively (Supplementary Figs. 5b and 11c), implying that oysters could be a vector of avian viruses for oysterivorus birds. Although we did not detect well-known foodborne viruses associated with oysters, such as AiV (*Kobuvirus*, *Picornaviridae*)^22,82^, HAV (*Hepatovirus*, *Picornaviridae*)^21,83–86^, or HEV (*Hepevirus*, *Hepeviridae*)^86–89^, viruses were detected from related families (*Picornaviridae* and *Hepeviridae*), including tentative zumaviruses (distantly related to AiV and HAV) and monhepeviruses (distantly related to HEV) (Supplementary Figs. 8a and 10a). Collectively, these findings reinforce the role of oysters as efficient natural accumulators of marine viruses and suggest their potential to serve as sentinels of vertebrate infecting viral contamination.

The presence of such phylogenetically diverse RNA viruses in oysters is explained by high-throughput filtration coupled with viral persistence. Adult oysters typically filter >100 L of seawater daily^4,5^, and in the process accumulate free viral particles, virus-infected cells, and virus-associated plankton. As oysters rely primarily on innate immunity^90–92^, ingested viruses, including vertebrate-associated viruses such as noroviruses can persist^71–75,93^. Thus, oysters can house phylogenetically diverse RNA viruses with broad probable host associations, resulting in a broad suite of environmental viruses rather than a subset of oyster-infecting viruses.

The ORV also provides the first synthesis on the distribution of diverse RNA virus assemblages across multiple oyster species, spanning eight countries on four continents (Figs. 1 and 4; Supplementary Figs. 1 and 3). Despite the wide geographic distribution, RNA virus community composition differed significantly among oyster species (PERMANOVA: r^2^ = 0.343, *p* = 0.001) and across countries (PERMANOVA: r^2^ = 0.520, *p* = 0.001), indicating strong oyster-species- and location-specific structuring of oyster viromes. Such patterns likely reflect differences in oyster microbiomes, diets, associated fauna, and local environmental conditions, suggesting that oyster viromes may provide informative snapshots of local viral landscapes.

Nonetheless, some groups of viruses were widely distributed. For example, vOTUs belonging to the *Calicividae* (e.g., ORV222), *Neornaviridae* (e.g., ORV319, ORV375, ORV386, and ORV448), and *Marnaviridae* (e.g., ORV225) were detected at >85% of the locations and in eight of the nine oyster species with detectable RNA viruses (Fig. 4). This pattern was particularly evident in Pacific oysters from British Columbia (BC), Canada, where viruses from these three families were detected in nearly all 96 metatranscriptomic samples collected from 13 locations, encompassing both farmed and wild oysters^38^ (Supplementary Fig. 13). Their broad distribution across oysters may reflect different aspects of oyster biology, with viruses in the *Marnaviridae* being consumed directly or in infected phytoplankton, the *Neornaviridae* representing viruses of oyster-associated symbionts, and viruses in the *Calicividae* being accumulated from surrounding seawater. Thus, despite strong geographic and host-associated differentiation across the overall virome, a small subset of RNA viruses (∼1% of total vOTUs) were broadly distributed across oysters and locations.

In addition, despite the substantial expansion of known oyster RNA virus diversity reported here, sampling remains strongly uneven among oyster species (Extended Data Table 1; Figs. 1 and 3; Supplementary Figs. 1 and 3). Although 774 of the 938 metatranscriptomes analyzed had not been included in previous RNA virus surveys, most recovered viruses were still associated with Hongkong oysters and Pacific oysters, reflecting the disproportionate availability of sequencing data for these two species. In future, more balanced sampling across oyster species will is required to obtain a representative estimate of RNA virus diversity in oysters.

In summary, this study demonstrates that oysters harbour a remarkably diverse array of RNA viruses, including many previously undescribed viruses, most of which likely infect other organisms. The discovery of novel viral families, genera, and host associations highlights that oysters accumulate environmental RNA viruses, including vertebrate-infecting viruses. These findings are consistent with oysters providing an important ecosystem service by removing environmental viruses from coastal waters that could pose health risks to other animals. Moreover, the wide range of RNA viruses found in oysters that infect other animals suggests that oysters are sentinels of coastal viral communities and could be used as sensors of viral contamination. Further investigations are needed to determine whether oysters function primarily as sinks, or may also be sources for viruses circulating in coastal ecosystems.

## Methods

### Acquisition of metatranscriptomic data for analysis

To investigate the RNA viruses associated with different species of oysters from a wide range of locations, we analyzed 96 transcriptomes obtained from Pacific oysters in BC, Canada^38^, as well as publicly available (as of 30 September 2024) oyster transcriptomes in the National Center for Biotechnology Information (NCBI), European Nucleotide Archive (ENA), and National Genomics Data Center (NGDC). These were combined to create a dataset of 938 metatranscriptomes from 45 studies, spanning 13 oyster species across 66 locations in 14 countries and five continents (Extended Data Table 1), consisting of ∼3 Tb of sequence data. These metatranscriptomes were generated from various sample types (e.g., tissue, hemolymph, or whole body) obtained from oysters at different developmental stages (e.g., larvae, juvenile, or adult) and in different health states (e.g., healthy, diseased, or under stress).

The dataset included the Hongkong oyster (*Magallana hongkongensis*), Pacific oyster (*Magallana gigas*), Olympia oyster (*Ostrea lurida*), Eastern oyster (*Crassostrea virginica*), Iwagaki oyster (*Magallana nippona*), Pearl oyster (*Pinctada fucata, Pinctada margaritifera* and *Pinctada imbricata radiata*), Flat oyster (*Ostrea edulis*), Sydney Rock Oyster (*Saccostrea glomerata*), Australian black-lip oyster (*Saccostrea sp.* non-mordax lineage J), Portuguese oyster (a.k.a., Fujian oyster: *Magallana angulata*) and Mangrove oyster (*Crassostrea tulipa*). Most metatranscriptomes were derived from Pacific and Hongkong oysters, with relatively few from other species.

### Sequence assembly and viral sequence detection

Raw reads from RNASeq samples were first quality-checked with fastQC (https://github.com/s-andrews/FastQC). Adapters and low-quality reads were trimmed by Trimmomatic v.0.38^94^ using the following settings: EADING:3 TRAILING:3 SLIDINGWINDOW:4:15 MINLEN:36. These quality-controlled (QC) pair-end reads from each sample were *de novo* assembled using MEGAHIT v.1.2.9^95^ (parameters: --min-contig-len 1000 --k-min 21 --k-max 141). The contigs were subsequently dereplicated using the dedupe.sh script from BBTools (https://jgi.doe.gov/data-and-tools/software-tools/bbtools/). Then, contigs <1 kb were removed using Seqtk v.1.3 (parameters: seq -L 1000; https://github.com/lh3/seqtk). Finally, this generated 10,373,790 deduplicated contigs (>1kb) from all samples for the analyses outlined below.

To screen the contigs belonging to RNA viruses, the RdRp-scan pipeline^96^ (https://github.com/JustineCharon/RdRp-scan) with default settings was employed. This pipeline uses RdRp-specific hidden Markov models (HMMs) and structural comparisons to efficiently detect and taxonomically annotate divergent RNA viruses in metatranscriptomic datasets based on the detection of RdRp sequences. Prior to the analysis, gene-prediction with ORFfinder v.1.8 (https://www.ncbi.nlm.nih.gov/orffinder/) was applied to the >1-kb contigs using gene code 1 to 33. Genes predicted by each gene code were combined and deduplicated using the dedupe.sh script from BBTools (https://jgi.doe.gov/data-and-tools/software-tools/bbtools/). Then, the deduplicated gene fragments were searched for RdRp motifs using HMMER v.3.3.2^97^. The genes containing RdRp motifs were kept, and their original contigs extracted using Seqkit v.2.2.0^98^. In addition, RdRp-containing contigs were obtained using an alternative BLAST approach, in which BLASTx was used to screen the obtained >1-kb contigs sequences against the ’RdRp-scan_0.90’ database v.2.0.9^96^ using an e-value cutoff of 10^-5^ and Diamond v.2.0.15 (https://github.com/bbuchfink/diamond) with the following parameters: -d RdRp-scan_0.90.dmnd --very-sensitive --matrix BLOSUM45 -e 1e-5 --min-orf 600 --max-target-seqs 1. Positive hits from both the HMM-search and Diamond-BLAST approaches were combined and considered as the RdRp-containing contigs. These contigs were deduplicated using the dedupe.sh script from BBTools (https://jgi.doe.gov/data-and-tools/software-tools/bbtools/), and completeness assessed using CheckV v.0.7.0^99^ with default parameters. Species-level viral populations (vOTUs) were clustered at 90% average nucleotide identity (ANI) across >75% of the alignment fragment (AF) using CheckV scripts^99^ (https://bitbucket.org/berkeleylab/checkv/src/master/).

To obtain the RdRp palm sequences, RdRp-containing contigs >1 kb were analyzed using palm_annot.py (https://github.com/rcedgar/palm_annot)^51,100^ with default parameters.

### Taxonomic assignment of ORV vOTUs

Taxonomic assignments of the >1 kb RdRp-containing contigs (ORV vOTUs were clustered using >90% ANI across >75% AF) was determined by leveraging the power of four complementary virus classification pipelines. First, the RdRp-scan pipeline^96^ was used again for primary classification. This involved using BLASTx with Diamond v.2.0.15^101^ to compare ORV vOTUs with the ’RdRp-scan_0.90’ database using an e-value cutoff of 10^-5^ to ascertain the best hit for each vOTU (parameters: -d RdRp-scan_0.90.dmnd --very-sensitive --matrix BLOSUM45 -e 1e-5 --min-orf 600 --max-target-seqs 1). Second, RdRp-palm-based classification^51^ was performed by searching RdRp palm sequences derived from each ORV vOTU against PALMdb (released 26 April 2023; https://github.com/ababaian/palmdb) using Diamond v.2.0.15^101^ (BLASTp) with an e-value cutoff of 10^-5^ to identify the closest reference sequence (parameters: -d otu_centroids.fa.dmnd --ultra-sensitive --masking 0 -e 1e-5 --max-target-seqs 1). Third, ORV vOTUs were further classified employing the Kaiju pipeline v.1.9.1^102^ with the NCBI nr database using default settings (parameters: -t nodes.dmp -f kaiju_db_nr_euk.fmi -n names.dmp). Fourth, contigs were annotated using the Contig Annotation Tool (CAT) v4.6^103^ with default parameters.

The taxonomy of each ORV vOTU was assigned by integrating the taxonomic information obtained from the four approaches described above, following a hierarchical priority order of RdRp-scan, RdRp-palm-based classification, Kaiju, and CAT. When conflicting assignments occurred at lower taxonomic ranks (e.g., family level) but a consistent higher-level classification (e.g., order level) was supported across methods, the shared higher-level taxonomy was retained. Finally, taxonomic assignments of RNA viruses were further refined and curated using phylogenetic analyses of RdRp-containing polyprotein sequences and/or RdRp palm sequences within each viral phylum, class, order, and/or family, by comparison with reference sequences retrieved from the NCBI RefSeq virus database (release 216), as described below.

### Host prediction

Virus-host predictions were made based on similarity of the ORV vOTU-encoded RdRp sequences with viral hits in the ’RdRp-scan_0.90’ database (https://github.com/JustineCharon/RdRp-scan)^96^. When host information was unavailable in the ‘RdRp-scan_0.90’ database, it was sourced from NCBI taxonomy (https://ftp.ncbi.nih.gov/pub/taxonomy/) using the ’taxonkit name2taxid’ tool (https://github.com/shenwei356/taxonkit)^104^. If additional validation was needed, manual searches for the putative hosts were made with either NCBI accession numbers or virus species names, using NCBI taxonomy (https://ftp.ncbi.nih.gov/pub/taxonomy/), ICTV VMR (release 37; https://ictv.global/vmr) and Virus-Host DB^105^ (https://www.genome.jp/virushostdb) databases. Finally, the putative host was manually refined based on the phylogenetic placement of each RdRp-containing polyprotein, and whether its closest phylogenetic groups (genus, family, order, or class) belonged to viruses infecting plants, algae, fungi, protists, bacteria, vertebrates, or invertebrates.

### Virus annotation

Putative coding sequences (CDSs) of ORV vOTUs were identified using ORFfinder v.1.8 (parameters: gene code = 11 for prokaryotic viruses and gene code = 1 for eukaryotic viruses; https://www.ncbi.nlm.nih.gov/orffinder/) for fragments >600 bp, and analyzed in Geneious Prime® 2023.2.1. Annotation of these CDSs, including identification of conserved domains within the polyprotein, was done using NCBI Conserved Domain Search (NCBI CD-Search; https://www.ncbi.nlm.nih.gov/Structure/bwrpsb/bwrpsb.cgi) and InterProScan v.5.59-91.0^106^. The InterProScan pipeline integrates a range of protein databases, including Pfam, SUPERFAMILY, Conserved Domains Database (CDD), PANTHER, ProSiteProfiles, CATH-Gene3D, Simple Modular Architecture Research Tool (SMART), and TIGRFAMs, enabling a detailed exploration of the protein profiles and domain architectures inherent to the viruses.

### Phylogenic analysis of ORV vOTUs

Evolutionary relationships were inferred from phylogenetic analysis of the RdRp-containing ORV vOTUs polyprotein sequences with representative RNA viruses from NCBI RefSeq (release 216). The RdRp amino-acid sequences were aligned using MAFFT version 7^107^ with default parameters. Then, the multiple alignments were trimmed using Clipkit v.1.4.1^108^ (parameters: -m kpic-smart-gap); maximum-likelihood (ML) phylogenies were conducted using these multiple alignments. The construction of the ML phylogeny was executed using IQ-TREE v.2.2.0.3^109^ with using 1,000 bootstrap replicates (parameters: -m MFP -B 1000 -bnni). This analysis employed the best-fit substitution model and associated model of among-site rate variation determined and selected by IQ-TREE for each trimmed alignment.

To complement the RdRp-containing polyprotein phylogenies, an additional phylogenetic analysis was performed using conserved RdRp palm sequences. Briefly, RdRp palm sequences from oyster-associated viruses and representative RNA viruses from NCBI RefSeq (release 216) were identified using palm_annot.py (https://github.com/rcedgar/palm_annot)^51,100^ with default parameters, as described above. These palm sequences were then aligned using MAFFT v.7, trimmed with Clipkit v.1.4.1, and used to construct maximum-likelihood phylogenies with IQ-TREE v.2.2.0.3 employing the same model selection and bootstrap settings as above.

### Comparative genomic analysis of RNA viruses

To assess whether there is species-level genome similarity (>90% ANI across >75% AF) among ORV vOTUs and RNA viruses from different hosts or habitats catalogued in the ICTV (release 37) and NCBI RefSeq (release 216) databases, as well as those identified from publicly available RNA viromic databases (e.g., Serratus^51^, RVMT^52^, Tara Ocean^54^, IMG/VR4^53^, and LucaProt^55^), we conducted a clustering analysis using all-versus-all BLAST (parameters: blastn -max_target_seqs 10000 -perc_identity 90). This analysis, which applied a threshold of >90% average nucleotide identity (ANI) across >75% alignment fraction (AF), was performed using the CheckV script^99^ (https://bitbucket.org/berkeleylab/checkv/src/master/). The vOTUs that did not share species-level similarity with any RNA virus in these databases (>90% ANI across >75% AF) were identified as novel viruses.

Putative new genera or families are proposed for vOTUs that form distinct, well-supported (bootstrap values >50%) phylogenetic clades with at least one novel virus based on the RdRp-containing polyprotein and/or RdRp palm sequences. Potential new families have <40% amino-acid identity (AAI) in the RdRp palm sequences^51,100^ when compared to members of neighbouring families. Similarly, potential new genera have <75% AAI in the RdRp palm sequences^51,100^ when compared to members of neighbouring genera.

### Profiling of abundances of RNA viruses detected in transcriptomes

The ‘abundance’ of ORV vOTUs was analyzed across oyster samples (each linked to a metatranscriptome with an NCBI/ENA/NGDC accession). Briefly, quality-filtered FASTQ files were mapped back to the ORV vOTUs sequences with an ANI >98%, using BBmap v.38.86^110^ (parameter: nodisk pigz=t usejni=t minid=0.98). The sequencing coverage or depth of each vOTU was calculated using the jgi_summarize_bam_contig_depths script from the MetaBAT2 pipeline v.2.15^111^ (parameters: minContigLength 36 --minContigDepth 2). For each analyzed metatranscriptomic sample, RPKM (reads per kilobase per million) normalization of mapped read counts was performed using CoverM v.0.6.1 (https://github.com/wwood/CoverM) with default parameters. This RPKM normalized abundance allows a quantitative assessment of ORV vOTUs distribution, facilitating comparisons across various oyster samples and revealing the relative levels of viral loads in the host.

### Statistical analyses

The community composition dissimilarity of RNA viruses across oyster species, developmental stages, sample types, health status, and countries of origin, was examined using fractional PERMANOVA^112^ with the adonis function and Bray-Curtis method in R^113^ package Vegan-v2.5^114^. The pair-wise post-hoc test was performed using the R package pairwise-Adonis-v0.4^115^.

## Data availability statement

Sequences of oyster-associated RNA viruses (ORV) and other supplementary materials, including trees and alignments, can be found in figshare^116^ via this link: https://doi.org:10.6084/m9.figshare.30907478. PONV1 sequences were deposited in GenBank in NCBI under accession numbers PQ030832 and PV579160^38^. Other ORV vOTU sequences assembled from Pacific oyster samples in BC, Canada were deposited in the European Nucleotide Archive (ENA) under accession numbers OZ557033 to OZ557222. The remaining ORV vOTU sequences were deposited in GenBase in NGDC under accession numbers C_AA502500.1 to C_AA502973.1, and in GenBank in NCBI under accession numbers BK083572 to BK083672.

## Acknowledgments

We are extremely grateful for the discussions with members of the Suttle lab on the analysis, interpretation, and presentation of the data. We thank the UBC Advanced Research Computing (ARC) facilities for their support in high-performance computing and data management. This work was generously supported by an award from the Gordon and Betty Moore Foundation (GBMF #5600). Additional support was provided by the Natural Sciences and Engineering Research Council of Canada (Discovery Grant RPGIN-2020-06515), Canada Foundation for Innovation, and the British Columbia Knowledge Development Fund (Project #25412) to C.A.S.

## Author Contribution Statement

K.X.Z. conducted the bioinformatic and genomic analyses. K.X.Z. and C.A.S. contributed to the interpretation of the results. K.X.Z. wrote the manuscript with input from A.M.C. and C.A.S.

## Conflict of Interest

The authors declare no competing financial interests.

## Extended Data Table Legends

**Extended Data Table 1 Publicly available oyster transcriptomes used for this study.**

**Extended Data Table 2 Metadata for the virus-associated RNA viruses (ORV) catalog.**

**Extended Data Table 3 RPKM-normalized abundance of RNA viruses in oysters across various locations.** For each sampling location (see details in Extended Data Table 1), **t**he RPKM-normalized abundance was determined by mapping metatranscriptomic reads to the 766 ORV vOTU genome sequences with an average nucleotide similarity (ANI) >98% and then normalizing the mapped reads to RPKM (reads per kilobase per million). It provides a semi-quantitative assessment of the examined viruses present in oysters across various locations, revealing the relative levels of viral loads.

## Supplementary Information

### I. Supplementary Results

#### Putative new viral families and genera inferred from phylogenetic divergence

Based on phylogenetic analysis, over half (434/766; 56.7%) of the ORV vOTUs are sufficiently divergent from RNA viruses in the NCBI RefSeq (release 216) that putatively they represent three new families and 16 novel genera (Fig. 2). The proposed families are the *Ymiviridae* in the order *Bunyavirales* (Supplementary Fig. 4a), *Megarnaviridae* in the order *Nidovirales* (Supplementary Fig. 5a), and the *Neornaviridae* (*Picobirnaviridae*-like) in the order *Durnavirales* (Supplementary Fig. 6a). The proposed genera are *Zinmovirus* in the family *Dicistroviridae* (order: *Picornavirales*, Supplementary Fig. 7a), *Zumavirus* in the family *Picornaviridae* (order: *Picornavirales*, Supplementary Fig. 8a), *Cedamavirus* and *Morcamavirus* in the family *Marnaviridae* (order: *Picornavirales*, Supplementary Fig. 7b), *Zipavirus* in the family *Secoviridae* (order: *Picornavirales*, Supplementary Fig. 7c), *Ravenavirus* in the family *Yueviridae* (order: *Goujianvirales*, Supplementary Fig. 4b), *Oystrovirus* in the family *Astroviridae* (order: *Stellavirales*, Supplementary Fig. 5a), *Gammanodavirus* in the family *Nodaviridae* (order: *Nodamuvirales*, Supplementary Fig. 9b), *Monhepevirus* in the family *Hepeviridae* (order: *Hepelivirales*, Supplementary Fig. 10a), *Monbirnavirus* in the family *Birnaviridae* (Supplementary Fig. 11a), *Kitornavirus* in the family *Avelrnaviridae* (order: *Sobelivirales*, Supplementary Fig. 5c), *Canulavirus* in the family *Tombusviridae* (order: *Tolivirales*, Supplementary Fig. 10b), *Tetramitovirus* in the family *Mitoviridae* (order: *Cryppavirales*, Supplementary Fig. 12a), *Mopavirus* and *Molavirus* in the family *Fiersviridae* (order: *Norzivirales*, Supplementary Fig. 12b), and *Moyavirus* in the family *Duinviridae* (order: *Norzivirales*, Supplementary Fig. 12b). These viruses occupy phylogenetic positions between families or genera (e.g., tentative *Ymiviridae*, *Neornaviridae*, *Oystrovirus*, *Cedamavirus*, *Zipavirus*, *Zumavirus*, *Monhepevirus*, *Canulavirus*, *Tetramitovirus*, *Mopavirus*, and *Moyavirus*), or are on deep branches within known families or genera (e.g., tentative *Megarnaviridae*, *Ravenavirus*, *Kitornavirus*, *Zinmovirus*, *Morcamavirus*, *Gammanodavirus*, *Monbirnavirus*, and *Molavirus*), enriching our understanding of RNA virus evolutionary diversity (Fig. 2).

#### Additional invertebrate-infecting RNA viruses in oysters

RNA viruses linked to oyster-associated invertebrates were also detected. ORV314, identified in Pacific oysters from France^1^ (Fig. 4), is closely related to the flatworm-infecting virus PSCNV^2^ within the family *Mononiviridae* (order *Nidovirales*; Supplementary Fig. 5a), suggesting a likely origin from worms inhabiting oysters^3–6^. Another *Nidovirales*-vOTU, ORV315, belonging to the family *Euroniviridae*, showed close phylogenetic proximity to Beihai nido-like virus 2 found in crabs (Supplementary Fig. 5a). ORV315 was detected in healthy oysters collected from naturalized beaches in BC (Supplementary Fig. 13), indicating that it may have been acquired from arthropods or arthropod-derived virus particles in the seawater, consumed as food by the oysters, or from arthropods associated with the oysters.

Additional invertebrate-infecting RNA viruses were also present, though it is unclear whether their hosts are oysters, oyster-associated invertebrates, or other environmental invertebrates. These include (+)ssRNA viruses from the *Sinhaliviridae* and *Nodaviridae* families within the order *Nodamuvirales* (Supplementary Fig. 9b), and *Dicistroviridae* in the order *Picornavirales* (Supplementary Fig. 7a), as well as (-)ssRNA viruses from the family *Yueviridae* in the order *Goujianvirales* (Supplementary Fig. 4b). For instance, three vOTUs (ORV481-483) assigned to the newly proposed *Ravenavirus* genus in the *Yueviridae* family formed a deep branch with *Yuyuevirus* members, such as Beihai sesamid crab virus 3 (Supplementary Fig. 4b). These negative-sense RNA viruses (*Ravenavirus*) were detected exclusively in Pacific oysters from BC, Canada, occurring in both wild (healthy) and farmed (diseased) populations, but not in Pacific oysters from other locations or in other oyster species (Fig. 4; Supplementary Fig. 13). This geographic and host specificity suggests an ecological or evolutionary relationship between *Ravenavirus* and Pacific oysters in BC.

As well, BC Pacific oysters harbor a sinaivirus (ORV545; *Sinhaliviridae* in the order *Nodamuvirales*) that clusters with honey-bees-infecting sinaiviruses^7^ (Supplementary Fig. 9b). ORV545 represents a novel sequence detected in only 3 of 46 diseased and dying farmed adult Pacific oysters in BC and was not observed in oysters from other regions worldwide (Fig. 4; Supplementary Fig. 13).

A total of 18 *Nodaviridae*-like vOTUs were identified within the order *Nodamuvirales*. Several, including complete genomes (e.g., ORV551), formed a distinct lineage tentatively designated *Gammanodavirus*, branching near insect-infecting alphanodaviruses but distant from fish-infecting betanodaviruses (Supplementary Fig. 9b), suggesting their invertebrate-infecting nature. These *Gammanodavirus* sequences were assembled from Pacific, Hongkong, and Olympia oysters, with close relatives (determined by mapping reads to genomes with ANI >98%) distributed broadly across oyster species (e.g., Iwagaki oysters) and locations (Fig. 4).

*Dicistroviridae*-like viruses showed similarly broad distribution, occurring in seven out of 13 oyster species (Fig. 4). These viruses, tentatively named *Zinmovirus*, formed a distinct cluster near enteroviruses and aparaviruses (Supplementary Fig. 7a). Given that members of the *Dicistroviridae*^8^ primarily infect invertebrates, the probable hosts of these tentative zinmoviruses are likely invertebrates, although whether they infect oysters, oyster-associated invertebrates, or other invertebrates, is unknown. Notably, the widespread presence of zinmoviruses and gammanodaviruses across oyster species and locations worldwide likely suggests their long-term ecological and/or evolutionary associations with oysters.

#### Additional vertebrate-infecting RNA viruses in oysters

Other vertebrate-infecting RNA viruses (ORV1-3, ORV14-16, and ORV572) include the genera *Orthoreovirus* in the family *Spinareoviridae* (order: *Reovirales*, Supplementary Fig. 11c), *Mamastrovirus* and *Oystrovirus* (tentative) in the family *Astroviridae* (order: *Stellavirales*, Supplementary Fig. 5b), and *Zumavirus* (tentative) in the family *Picornaviridae* (order: *Picornavirales*, Supplementary Fig. 8a). Among these, ORV572, in the genus *Orthoreovirus* and family *Spinareoviridae* (order: *Reovirales*, Supplementary Fig. 11c), is distantly related to Avian orthoreovirus that infects chicken (NC_015127). Similarly, ORV1 is deeply branched with bird-infecting astroviruses (*Avastrovirus*), but sufficiently divergent to belong to a proposed new genus, *Oystrovirus*. In contrast, ORV2 and ORV3 clustered with mamastroviruses; astroviruses in the genus *Mamastrovirus* typically infect mammals (Supplementary Fig. 5b). Notably, these mamastroviruses (ORV2 and ORV3) and oystroviruses (ORV1) were only found in Pacific oysters in BC, Canada (Fig. 4; Supplementary Fig. 13), while the orthoreovirus (ORV572) was only detected in Hongkong oysters, hinting that vertebrate-infecting viruses may be regional in distribution.

The oyster-associated *Picornaviridae*-like vOTUs (ORV14-16), classified as *Zumavirus* (tentative), were assembled from Hongkong oysters, with ORV14 and ORV15 featuring new sequences that lack species-level similarity to any RNA viruses in current databases (Supplementary Fig. 8a). These zumaviruses were detected across multiple oyster species, including Hongkong, Pacific, and flat oysters, as well as in both diseased and wild Pacific oysters from BC, Canada (Fig. 4; Supplementary Fig. 13). Phylogenetically, zumaviruses form a distinct cluster that branches alongside other *Picornaviridae* genera, including *Kobuvirus* (which includes *Aichivirus*) and *Hepatovirus* (which includes Hepatitis A virus). Given that *Picornaviridae* are known to infect vertebrates only, we speculate the vertebrate-infecting nature of these identified zumaviruses.

#### RNA viruses with host undetermined

Finally, there are viruses that could not be assigned to a host (Fig. 3; Extended Data Table 2), including vOTUs assigned to *Neornaviridae* (tentative) in the order *Durnavirales* (Supplementary Fig. 6a), *Ymiviridae* (tentative) in the order *Bunyavirales* (Supplementary Fig. 4a), and *Monhepevirus* (tentative) in the family *Hepeviridae* within the order *Hepelivirales* (Supplementary Fig. 10a). A substantial portion (17.9% of ORV vOTUs; 137/766) of *Picobirnaviridae*-like vOTUs belonged to *Neornaviridae* (Fig. 3). These *Neornaviridae* vOTUs have *Picobirnaviridae*-like RdRp (Supplementary Fig. 6a) that were phylogenetically branched between the vertebrate-infecting *Picobirnaviridae* and fungi-infecting viruses of the order *Durnavirales* (e.g., *Curvulaviridae*, *Hypoviridae*, and *Fusariviridae*), a branching pattern likewise observed in previous studies^9,10^. Thus, the hosts of these *Neornaviridae* members are undetermined, although a recent study suggests that they likely infect microbes^11^. Notably, these *Neornaviridae* (*Picobirnaviridae*-like viruses) were widespread across all oyster species in almost all locations (45 out of 46) where RNA viruses were detected, as well as in all farmed and wild Pacific oysters in BC (Fig. 4; Supplementary Fig. 13).

Likewise, the host of *Bunyavirales*-like vOTUs (ORV484-486) was undetermined. Their RdRp-containing polyproteins clustered in one clade, sufficiently divergent to be a proposed new family, *Ymiviridae,* that branches closely with some families in the order *Bunyavirales*, including *Leishbuviridae*, *Phenuiviridae*, *Mypoviridae,* and *Arenaviridae*, but more distantly to members of the *Hantaviridae*, *Wupedeviridae,* and *Nairoviridae* (Supplementary Fig. 4a). These (-)ssRNA segmented viruses (*Ymiviridae*) include three vOTUs from both diseased Pacific oysters from farms and healthy naturalized beach oysters in BC and Flat oysters in France (Fig. 4; Supplementary Fig. 13).

Similarly, three *Hepeviridae*-like (hepe-like) viruses have been detected in global oyster populations, with two (ORV564 and ORV565) assembled from Hongkong oysters from China and one with a complete genome (ORV566) from Pacific oysters in BC, Canada (Fig. 4; Supplementary Figs 10a and 13). Phylogenetic analysis of the RdRp-containing polyprotein shows that these oyster-associated hepe-like viruses form a distinct clade, branching between the hepe-like virus detected in the invertebrate white-backed planthopper (*Sogatella furcifera*)^12^ and the known vertebrate-infecting members of *Hepeviridae*, including rana hepevirus, avihepevirus, chirohepevirus, rocahepevirus, and piscihepevirus (Supplementary Fig. 10a). Based on genetic divergence, these hepe-like viruses are proposed to constitute a new genus, *Monhepevirus* (tentative), within the family *Hepeviridae* (order *Hepevirales*). These hepe-like viruses were found in both diseased farmed oysters and healthy naturalized Pacific oysters in BC, as well as Hongkong oysters across multiple regions in China (Fig. 4; Supplementary Fig. 13). However, whether they infect invertebrates (e.g., oysters) or vertebrates is unknown.

### II. Supplementary Figures

**Supplementary Fig. 1.**
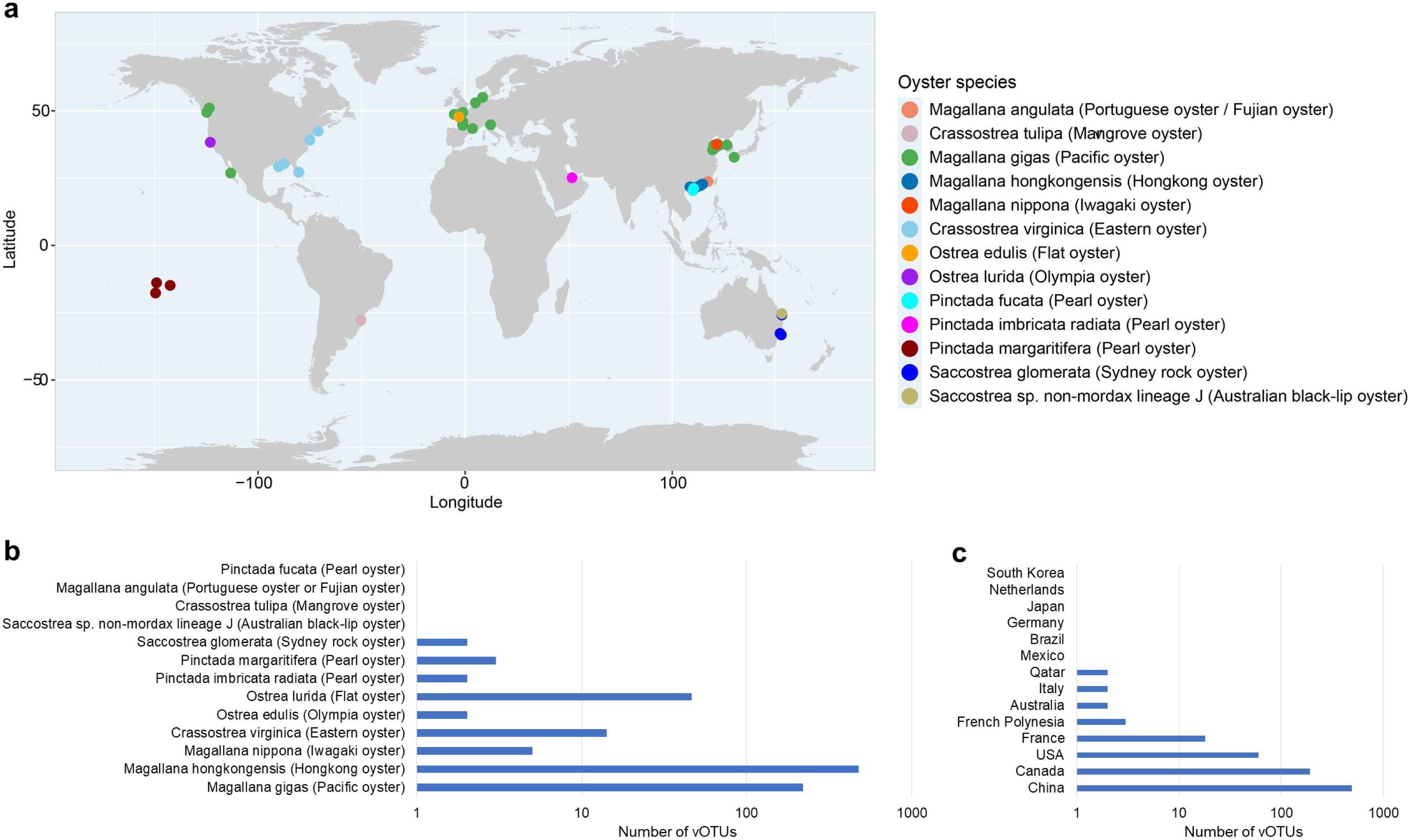
Overview of the oyster-associated RNA viruses (ORV) catalog. **a**, Map of sampling locations from which oyster transcriptomic samples were obtained (see details in Extended Data Table 1). Countries highlighted in black represent those from which transcriptomic samples of oysters were publicly available and have been examined in this study. **b**, Distribution of the number of viral operational taxonomic units (vOTUs) in the ORV catalog across oyster species. **c**, Distribution of the number of vOTUs in the ORV catalog across countries with available oyster samples.

**Supplementary Fig. 2.**
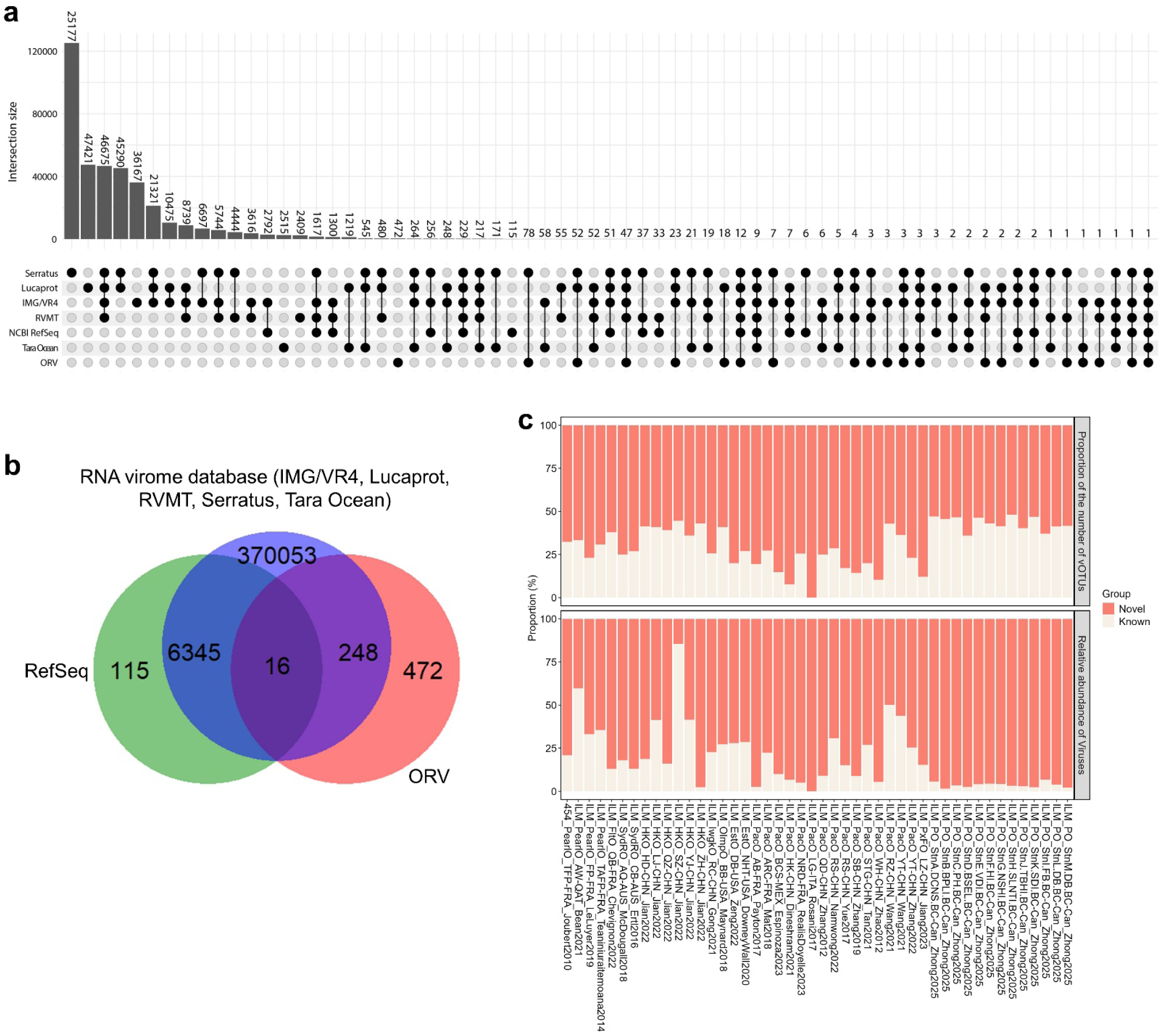
Specificity of RNA viruses found in global oyster populations. **a**, Genomic comparison of oyster RNA viruses (ORV) vOTUs with RNA viruses sequences found in the NCBI RefSeq (release 216) databases and the publicly available RNA virome databases such as Serratus^13^, Tara Ocean^14^, IMG/VR4^15^, RVMT^16^, and LucaProt^17^. Black dots indicate the presence of species-level similar viruses (>90% ANI across >75% AF) between two databases, while grey dots represent their absence. Black lines connecting black dots indicate the viruses detected across multiple datasets. The upper panel summarizes the total number of viruses for each black dot-line combination. A black dot without line connections, detected only in ORV catalog, indicates that the viruses are specific to oysters and do not share species-level similarity (>90% ANI across >75% AF) with RNA virus sequences in other datasets, suggesting possible specificity or novel viruses. **b**, Summary of novel RNA viruses detected in oysters. The Venn diagram illustrates the number of shared and unique species-level viral clusters (i.e., vOTUs) of RNA viruses among oysters (ORV catalog), the NCBI RefSeq (release 216), as well as the publicly available RNA virome databases such as Serratus^13^, Tara Ocean^14^, IMG/VR4^15^, RVMT^16^, and LucaProt^17^. **c**, Distribution of known and novel viral clusters (vOTUs) across sampling locations. The upper panel shows the proportion of the number of vOTUs detected at each location (n = 46), while the lower panel displays the relative abundance of these vOTUs. Locations where no RNA viruses were detected are excluded from the visualization (n = 20; see details in Extended Data Table 3).

**Supplementary Fig. 3.**
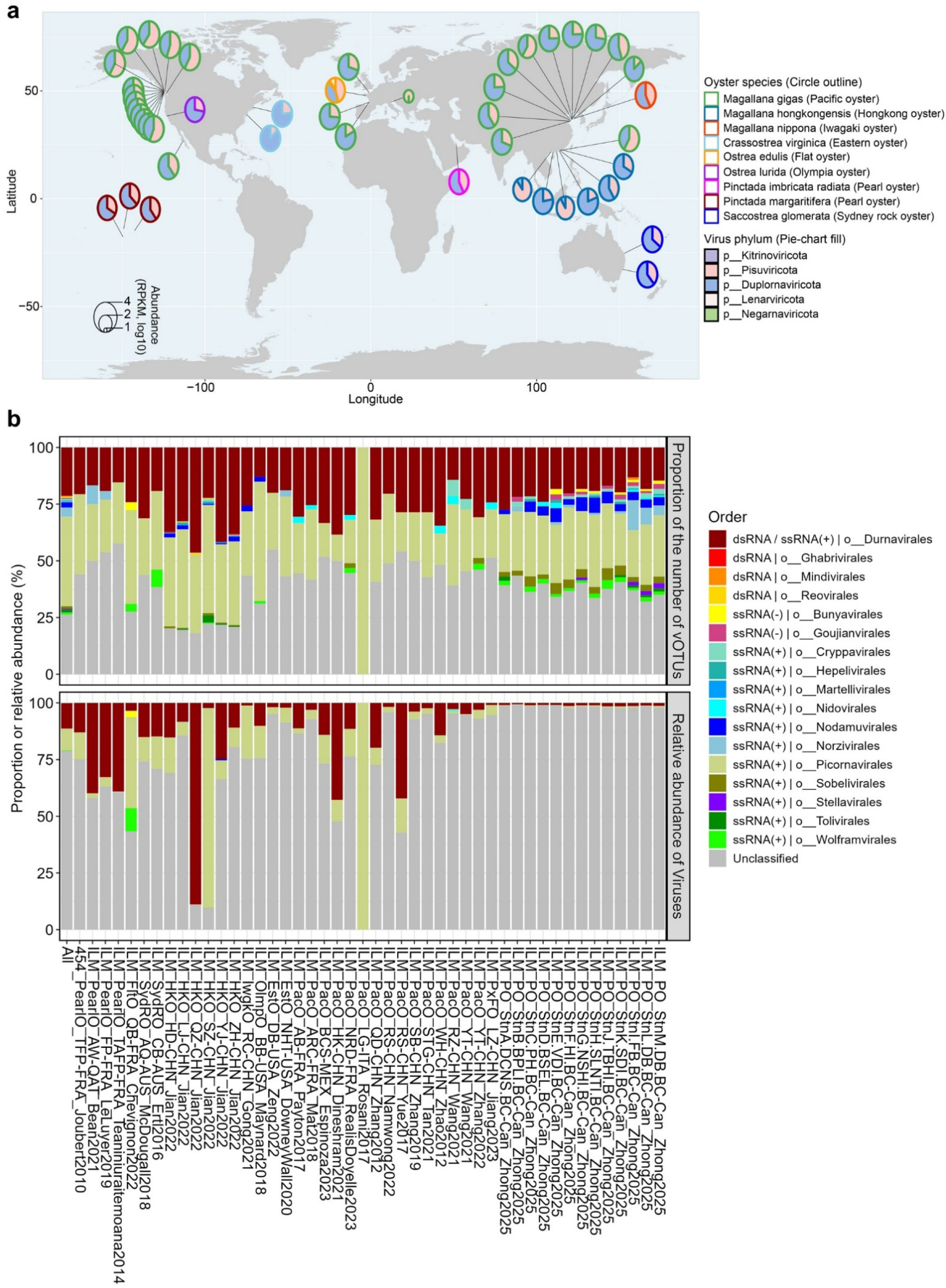
Taxonomic distribution oyster-associated RNA viruses (ORV) in oysters across locations. **a**, Phylum-level taxonomy distribution of the ORV vOTUs across locations where oyster transcriptomes were available. Each circle’s color specifies the oyster species, and its size reflects the normalized abundance of their RNA viruses, determined by the count of transcriptomic reads aligned to the detected ORV vOTUs genome (ANI >98%) in specific sites and adjusted by Reads Per Kilobase Million (RPKM). This RPKM normalized abundance allows a semi-quantitative assessment of RNA virus distribution, facilitating comparisons across species, studies, and/or locations. Oyster species showing a normalized abundance of zero are excluded from the visualization (n = 4; see details in Supplementary Fig. 1b and Extended Data Table 3). **b**, Order-level taxonomy distribution of vOTUs in oysters across sampled locations. The upper panel shows the proportion of the number of vOTUs detected at each location (n = 46), while the lower panel displays the relative abundance of these vOTUs. Locations where no RNA viruses were detected are excluded from the visualization (n = 20; see details in Extended Data Table 3). All indicate the combination of all locations with RNA viruses detected.

**Supplementary Fig. 4.**
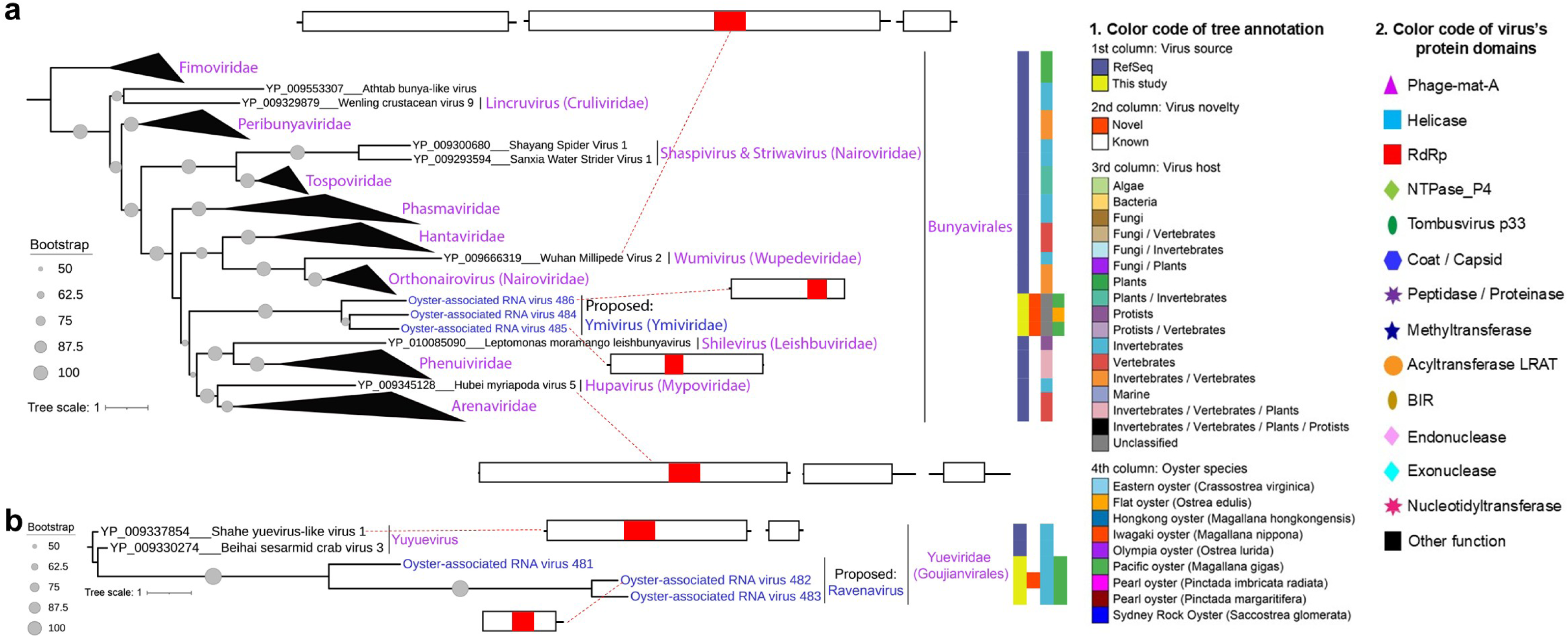
RdRp-containing polyprotein phylogeny of oyster-associated RNA viruses (ORV) in the orders *Bunyavirales* and *Goujianvirales* from phylum *Negarnaviricota*. **a**, Phylogeny of RNA viruses in the order *Bunyavirales*. The maximum-likelihood (ML) phylogenetic tree, based on the inferred amino acid sequence of the polyprotein containing RNA-dependent RNA polymerase (RdRp), illustrates the evolutionary relationships between ORV vOTUs (blue text at tree tips) and RNA viruses from the NCBI RefSeq (release 216; black text at tree tips). Node labels indicate bootstrap support values. Tree leaves are annotated to indicate virus source (1^st^ column; NCBI RefSeq vs. ORV vOTUs from this study), novelty (2^nd^ column; known vs. novel), derived oyster species (3^rd^ column), and predicted host groups (4^th^ column). The tree is grouped based on shared taxonomy, labeled in pink, and the proposed new taxonomy (genus or family) is labeled in blue when ORV vOTUs are sufficiently divergent and supported by strong bootstrap values. Representative virus genome map with annotated protein domains is displayed adjacent to the tree leaf, connected by a red dashed line. The ML tree is re-rooted using members of the family *Fimoviridae*. **b**, Phylogeny of RNA viruses in the order *Goujianvirales*. The ML phylogenetic tree is unrooted. Tree and virus annotations follow the same conventions as in panel a.

**Supplementary Fig. 5.**
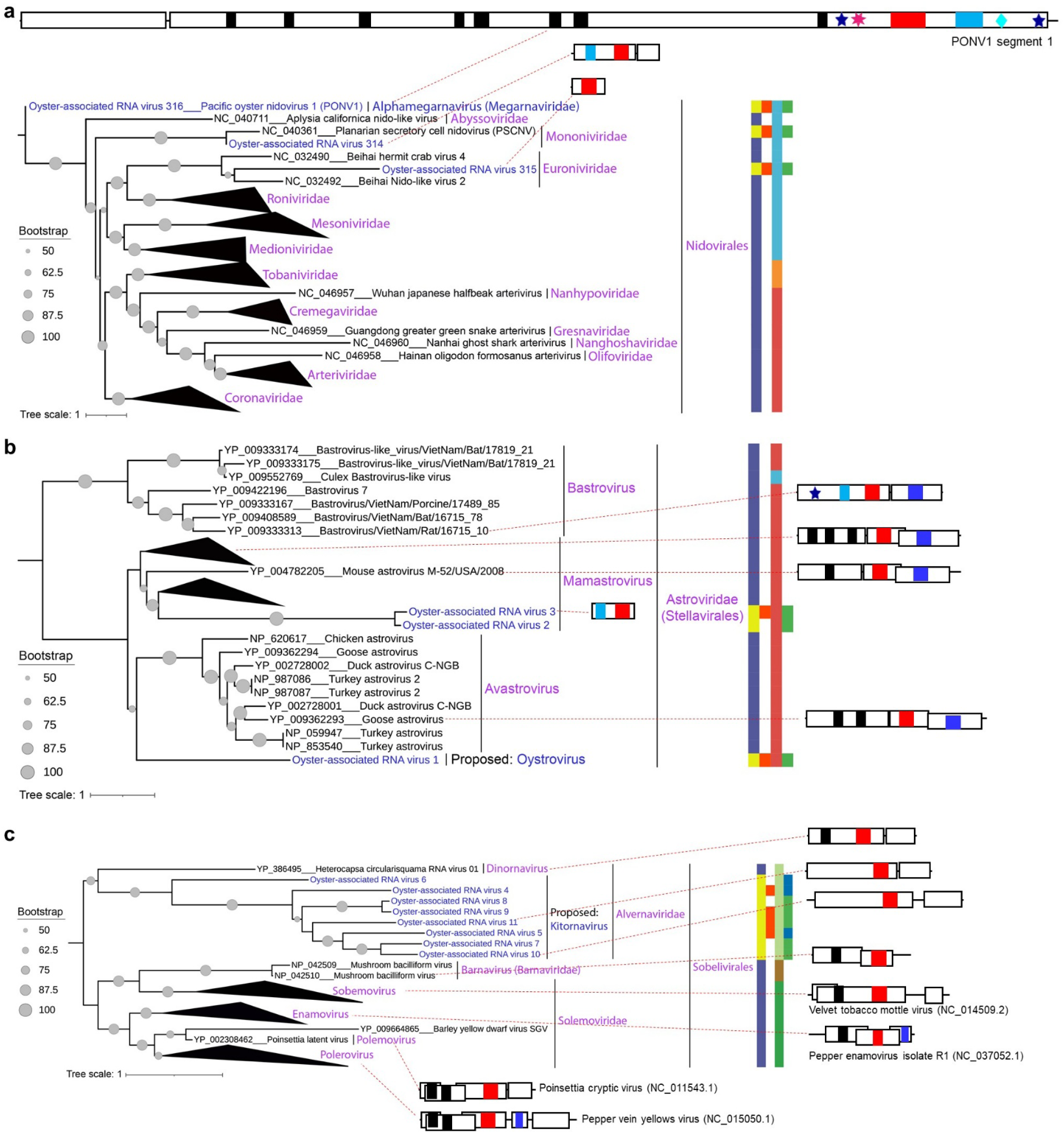
RdRp-containing polyprotein phylogeny of oyster-associated RNA viruses (ORV) in the orders *Nidovirales*, *Stellavirales*, and *Sobelivirales* from phylum *Pisuviricota*. **a**, Phylogeny of RNA viruses in the order *Nidovirales*. The ML phylogenetic tree is unrooted. **b**, Phylogeny of RNA viruses in the order *Stellavirales*. The maximum-likelihood (ML) phylogenetic tree is re-rooted using members of *Bastrovirus* from the family *Astroviridae*. **c**, Phylogeny of RNA viruses in the order *Sobelivirales*. The ML phylogenetic tree is re-rooted using members of the family *Alvernaviridae*. The legend for the figure (panels **a**, **b**, and **c**), including color key, tree annotations, and virus annotations, is the same as in Supplementary Fig. 4.

**Supplementary Fig. 6.**
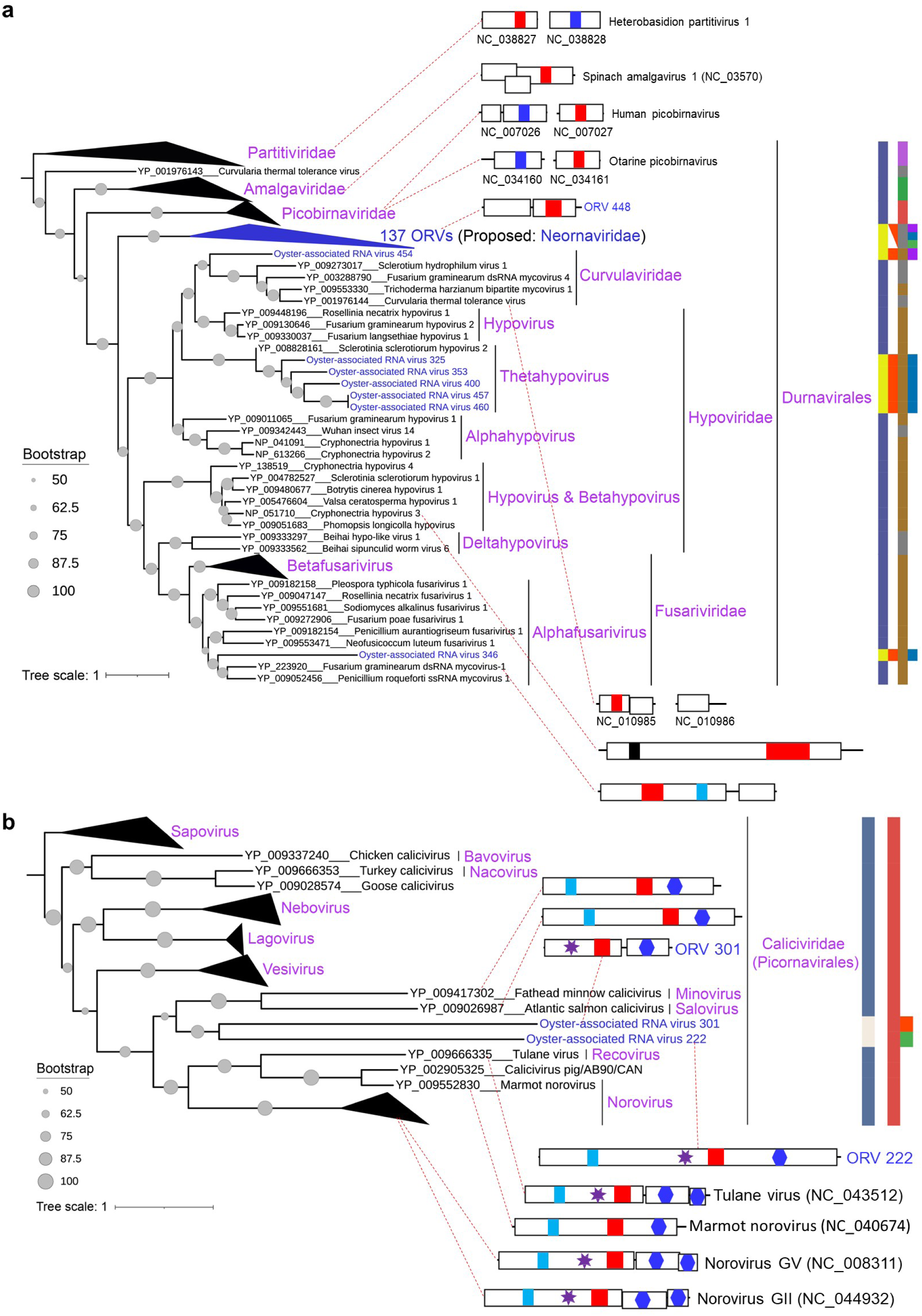
RdRp-containing polyprotein phylogeny of oyster-associated RNA viruses (ORV) in the order *Durnavirales* and the family *Caliciviridae* (order *Picornavirales*) from phylum *Pisuviricota*. **a**, Phylogeny of RNA viruses in the order *Durnavirales*. The maximum-likelihood (ML) phylogenetic tree is re-rooted using members of the family *Partitiviridae*. **b**, Phylogeny of RNA viruses in the family *Caliciviridae* from order *Picornavirales*. The ML phylogenetic tree is re-rooted using members of the genus *Sapovirus*. The legend for the figure (panels **a** and **b**), including color key, tree annotations, and virus annotations, is the same as in Supplementary Fig. 4.

**Supplementary Fig. 7.**
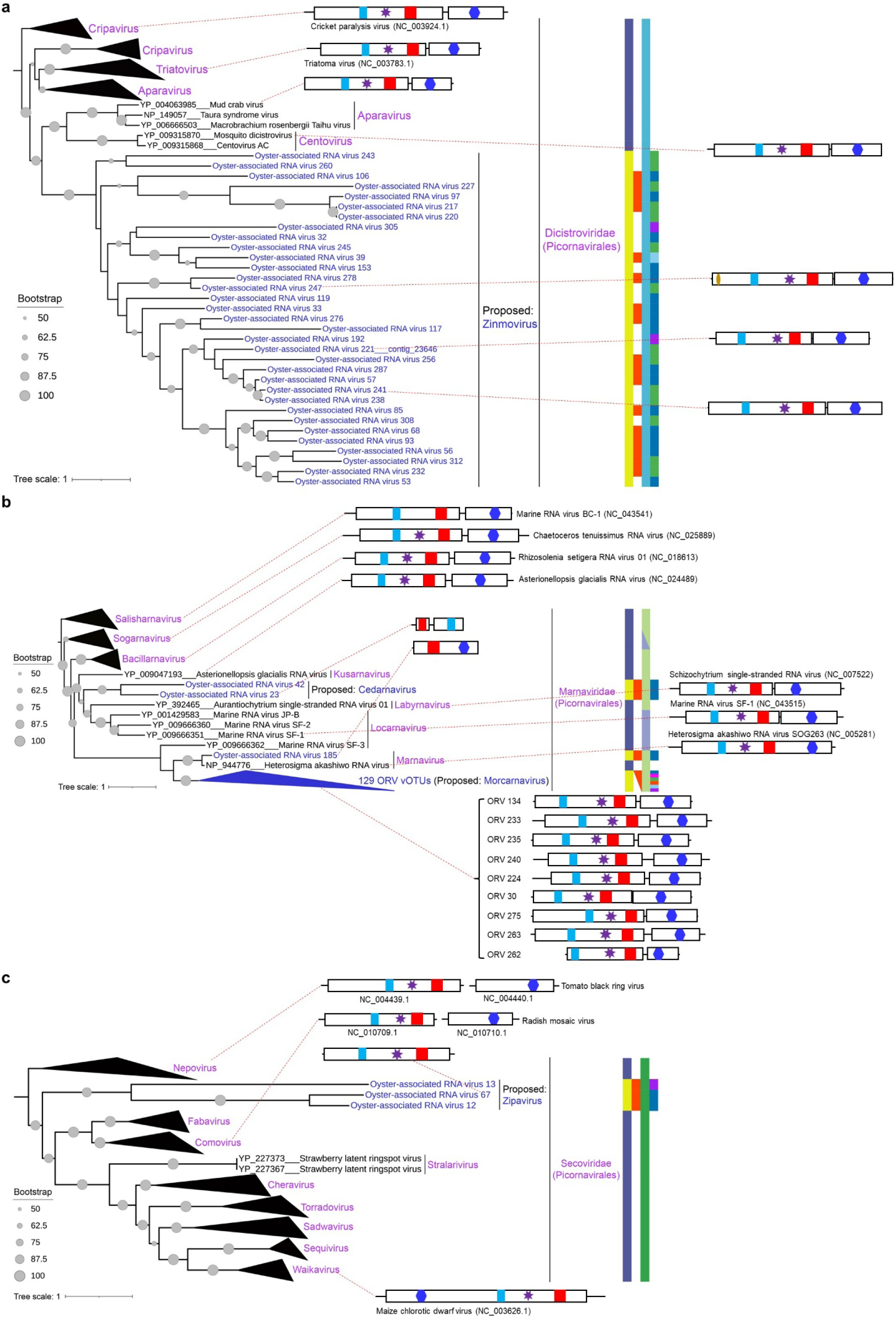
RdRp-containing polyprotein phylogeny of oyster-associated RNA viruses (ORV) in the families *Dicistroviridae, Marnaviridae, Secoviridae within the* order *Picornavirales* from phylum *Pisuviricota*. **a**, Phylogeny of RNA viruses in the family *Dicistroviridae* from order *Picornavirales*. The maximum-likelihood (ML) phylogenetic tree is re-rooted using members of the genus *Cripavirus*. **b**, Phylogeny of RNA viruses in the family *Marnaviridae* from order *Picornavirales*. The ML phylogenetic tree is re-rooted using members of the genus *Salisharnavirus*. **c**, Phylogeny of RNA viruses in the family *Secoviridae* from order *Picornavirales*. The ML phylogenetic tree is re-rooted using members of the genus *Nepovirus*. The legend for the figure (panels **a**, **b**, and **c**), including color key, tree annotations, and virus annotations, is the same as in Supplementary Fig. 4.

**Supplementary Fig. 8.**
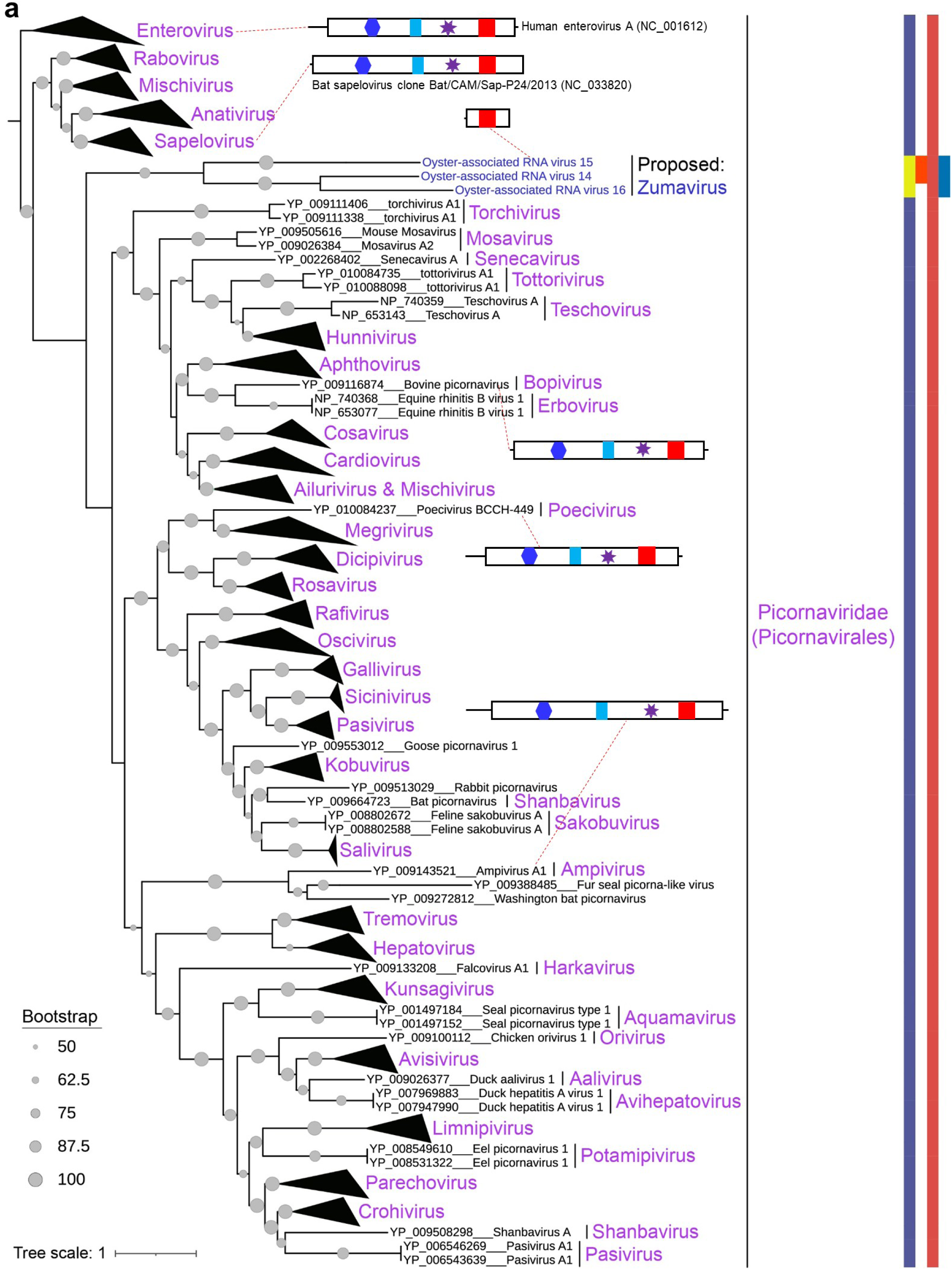
RdRp-containing polyprotein phylogeny of oyster-associated RNA viruses (ORV) in the family *Picornaviridae* within the order *Picornavirales* from phylum *Pisuviricota*. **a**, Phylogeny of RNA viruses in the family *Picornaviridae* from order *Picornavirales*. The ML phylogenetic tree is re-rooted using members of the genus *Enterovirus*. The legend for the figure, including color key, tree annotations, and virus annotations, is the same as in Supplementary Fig. 4.

**Supplementary Fig. 9.**
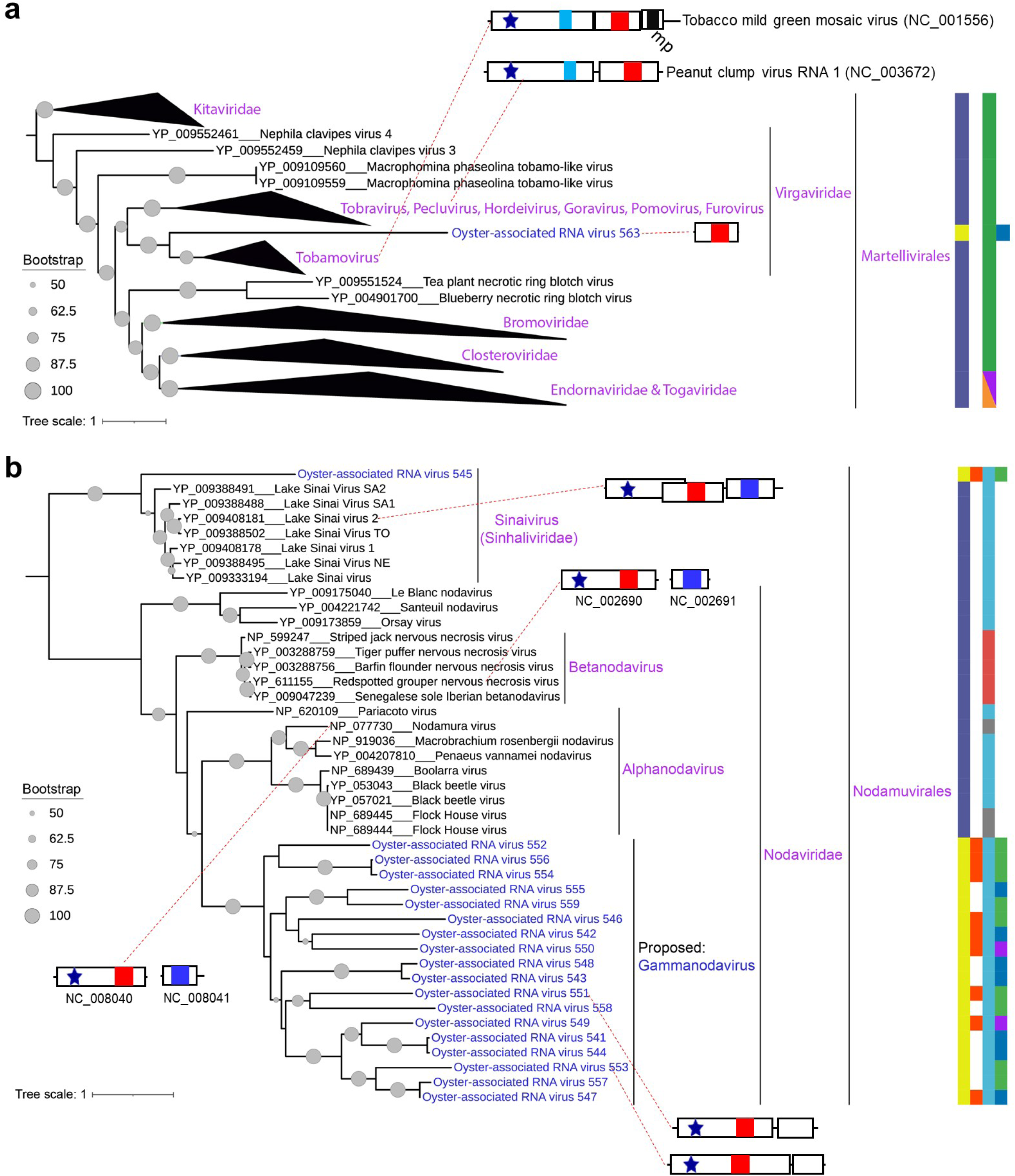
RdRp-containing polyprotein phylogeny of oyster-associated RNA viruses (ORV) in the orders *Martellivirales* and *Nodamuvirales* from phylum *Kitrinoviricota*. **a**, Phylogeny of RNA viruses in the order *Martellivirales*. The maximum-likelihood (ML) phylogenetic tree is re-rooted using members of the family *Kitaviridae*. **b**, Phylogeny of RNA viruses in the order *Nodamuvirales*. The ML phylogenetic tree is re-rooted using members of the family *Sinhaliviridae*. The legend for the figure (panels **a** and **b**), including color key, tree annotations, and virus annotations, is the same as in Supplementary Fig. 4.

**Supplementary Fig. 10.**
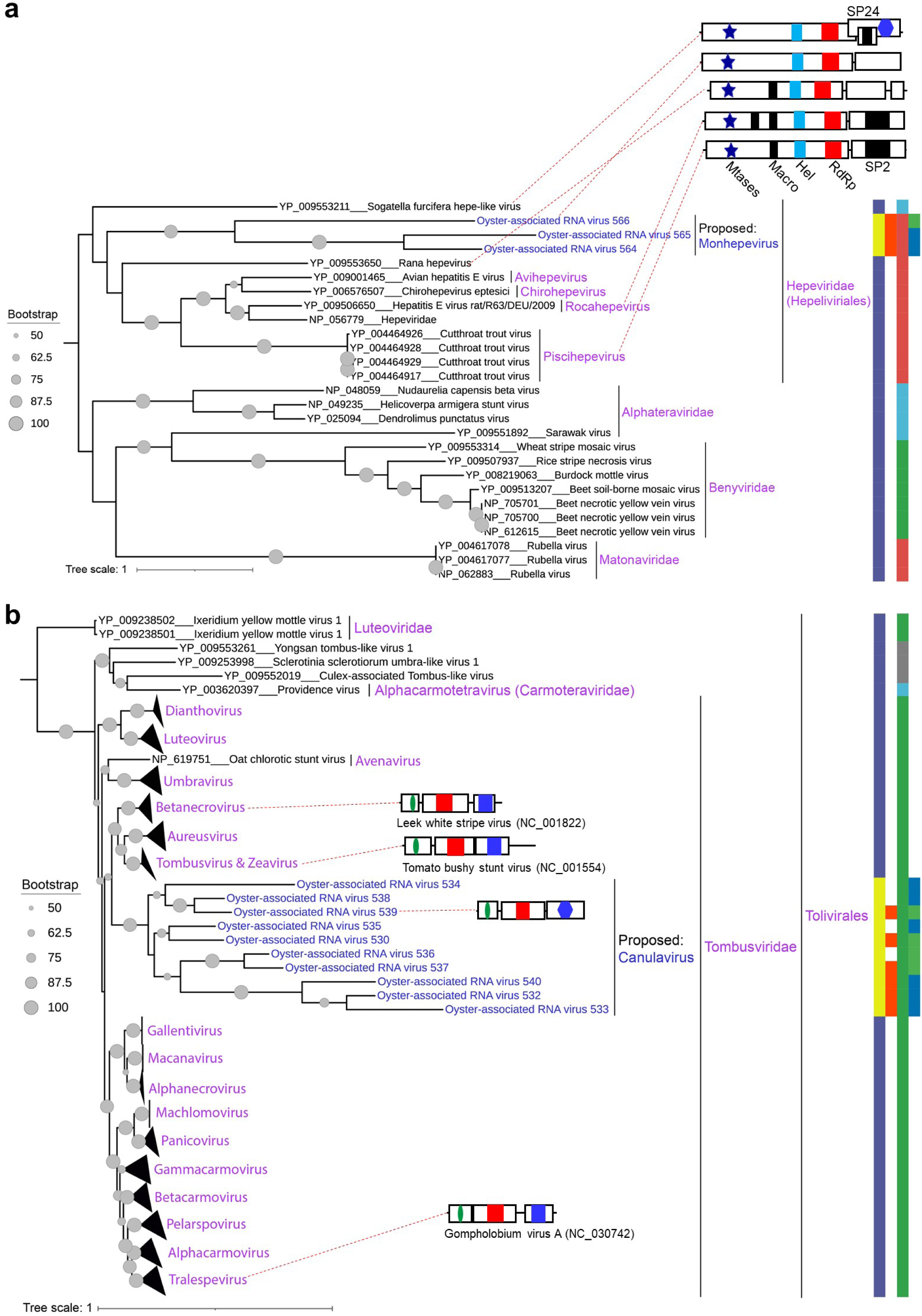
RdRp-containing polyprotein phylogeny of oyster-associated RNA viruses (ORV) in the orders *Hepelivirales* and *Tolivirales* from phylum *Kitrinoviricota*. **a**, Phylogeny of RNA viruses in the order *Hepelivirales*. The maximum-likelihood (ML) phylogenetic tree is re-rooted using members of the family *Hepeviridae*. **b**, Phylogeny of RNA viruses in the order *Tolivirales*. The ML phylogenetic tree is re-rooted using members of the family *Luteoviridae*. The legend for the figure (panels **a** and **b**), including color key, tree annotations, and virus annotations, is the same as in Supplementary Fig. 4.

**Supplementary Fig. 11.**
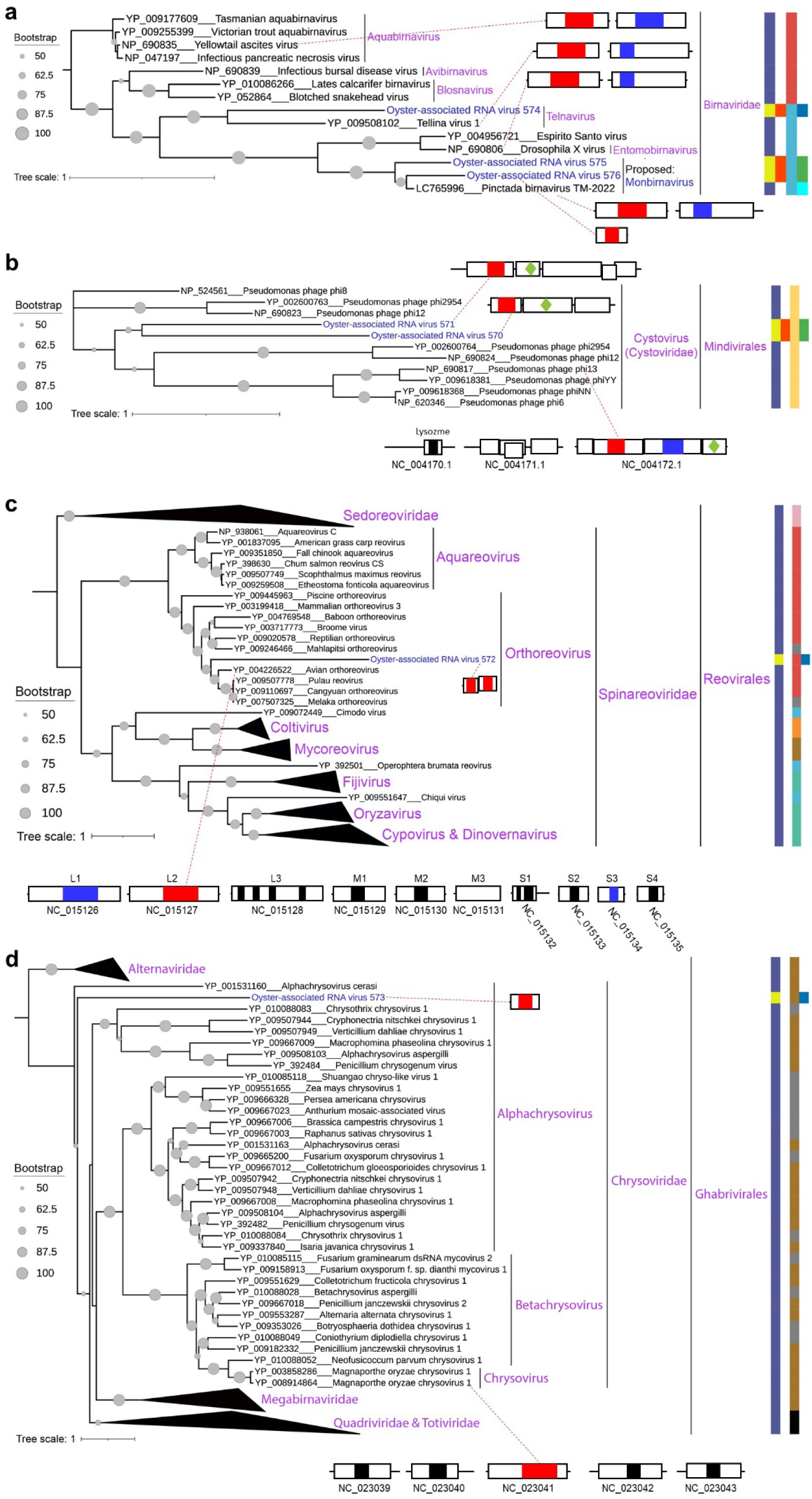
RdRp-containing polyprotein phylogeny of oyster-associated RNA viruses (ORV) in the family *Birnaviridae* and the orders *Mindivirales*, *Reovirales*, and *Ghabrivirales* from phylum *Duplornaviricota*. **a**, Phylogeny of RNA viruses in the family *Birnaviridae*. The maximum-likelihood (ML) phylogenetic tree is re-rooted using members of the genus *Aquabirnavirus*. **b**, Phylogeny of RNA viruses in the order *Mindivirales*. The ML phylogenetic tree is unrooted. **c**, Phylogeny of RNA viruses in the order *Reovirales*. The ML phylogenetic tree is re-rooted using members of the family *Sedoreoviridae*. **d**, Phylogeny of RNA viruses in the order *Ghabrivirales*. The ML phylogenetic tree is re-rooted using members of the family *Alternaviridae*. The legend for the figure (panels **a**, **b**, **c**, and **d**), including color key, tree annotations, and virus annotations, is the same as in Supplementary Fig. 4.

**Supplementary Fig. 12.**
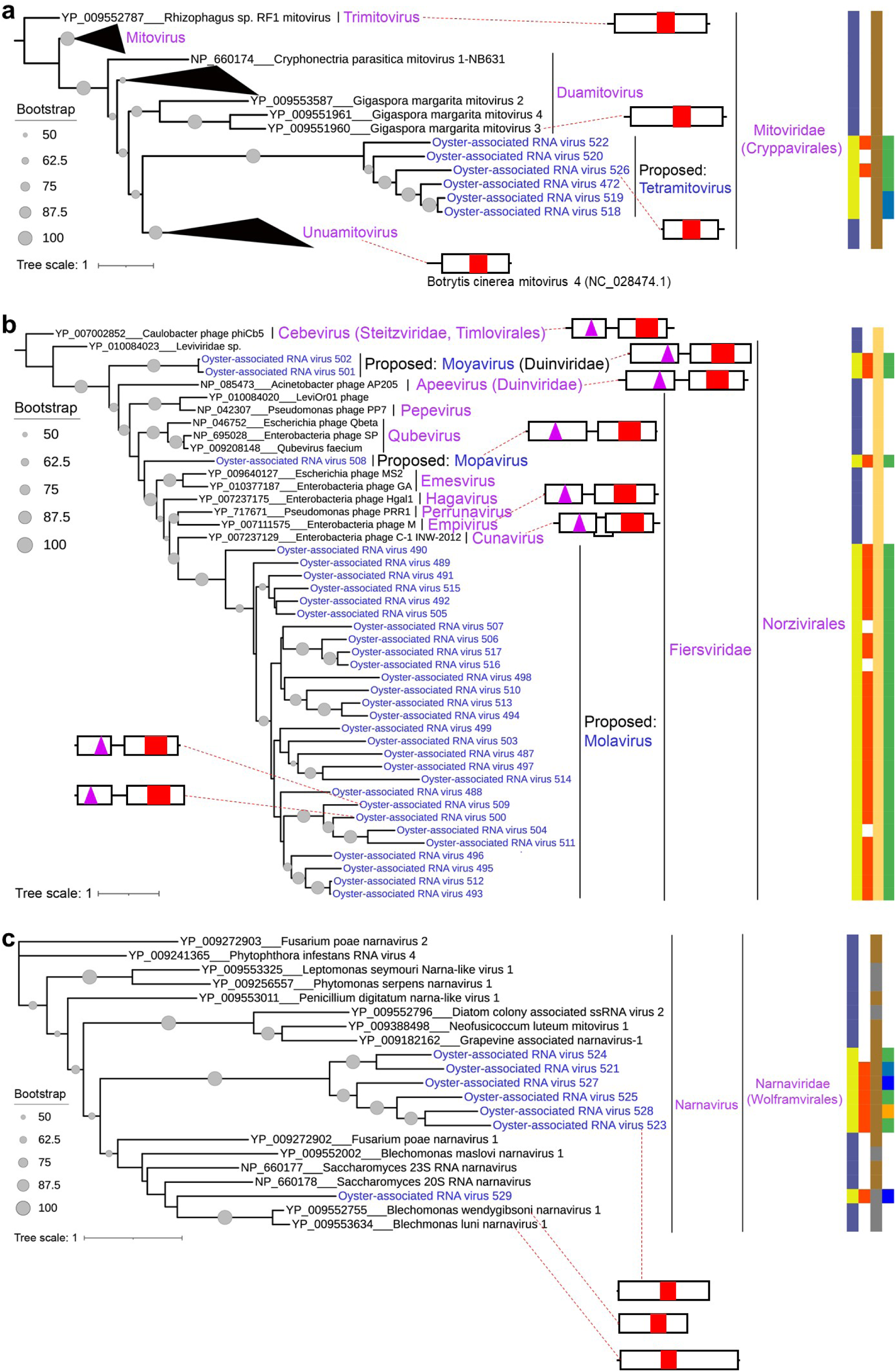
RdRp-containing polyprotein phylogeny of oyster-associated RNA viruses (ORV) in the orders *Cryppavirales* and *Norzivirales* from phylum *Lenarviricota*. **a**, Phylogeny of RNA viruses in the order *Cryppavirales*. The ML phylogenetic tree is re-rooted using members of *Trimitovirus* from the family *Mitoviridae*. **b**, Phylogeny of RNA viruses in the order *Norzivirales*. The ML phylogenetic tree is re-rooted using members of *Cebevirus* from the family *Steitzviridae* in the order *Timlovirales*. **c**, Phylogeny of RNA viruses in the order *Wolframvirales*. The ML phylogenetic tree is unrooted. The legend for the figure (panels **a**, **b**, and **c**), including color key, tree annotations, and virus annotations, is the same as in Supplementary Fig. 4.

**Supplementary Fig. 13.**
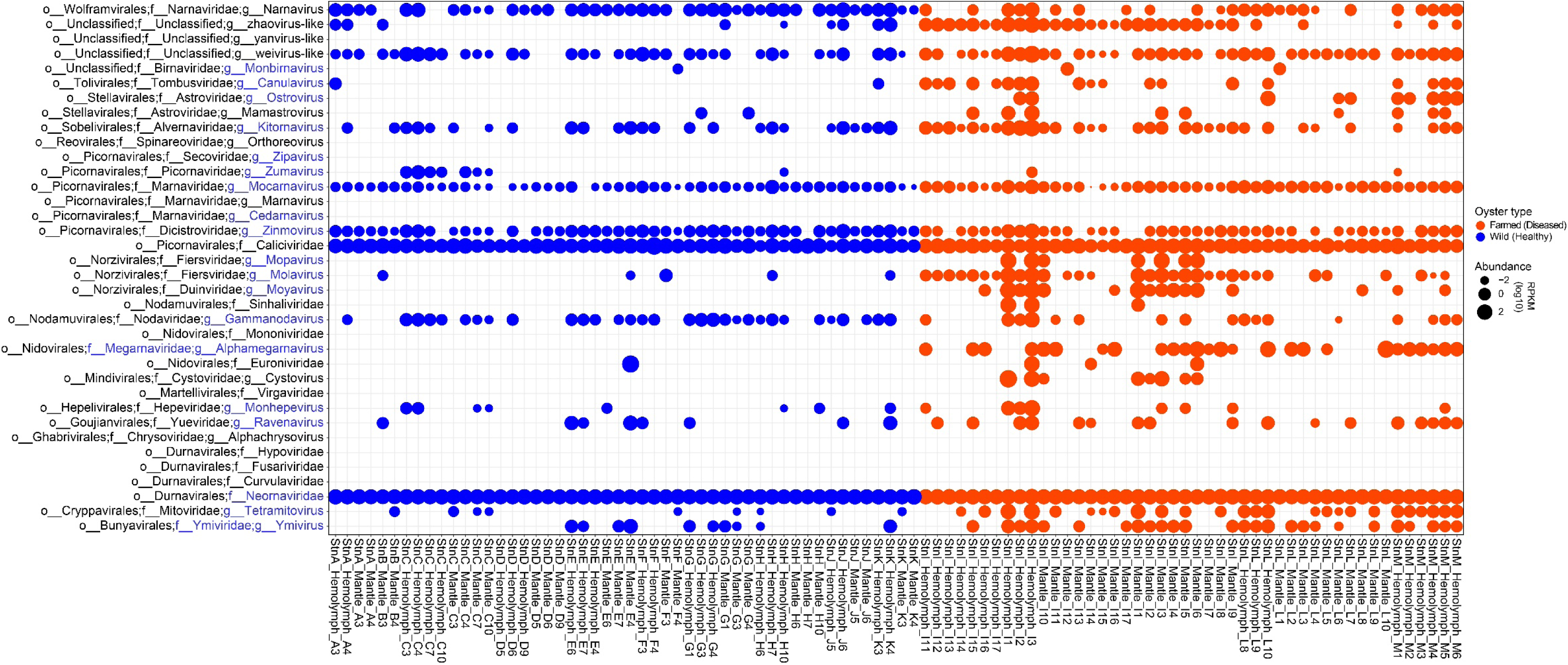
Prevalence of oyster-associated RNA viruses (ORV) assigned to newly proposed families and genera in adult Pacific oysters in British Columbia (BC), Canada. Each circle represents the detection of viruses, with the circle size indicating the RPKM-normalized abundance of these viruses in each oyster samples (n = 96). The RPKM (reads per kilobase pair per million) normalization involves mapping metatranscriptomic reads to ORV vOTUs of the viral families or genera with an average nucleotide similarity (ANI) of over 98%, providing a semi-quantitative measure of viral load across different oyster samples. The circle color distinguishes the wild (StnA, StnB, StnC, StnD, StnE, StnF, StnG, StnH, StnJ, and StnK) and farmed (StnI, StnL, and StnM) adult Pacific oysters sampled in BC, Canada. The farmed oysters were experiencing massive mortality events and were sampled from two farms [farm 1 (StnI) and farm 2 (StnL and StnM)] in Vancouver Island in the summer of 2020 (Extended Data Table 1). While the wild oysters were sampled from ten wild naturalized beach locations (StnA-H, StnJ, and StnK), the surrounding region of the farms experienced mortality incidents. Each oyster had its mantle tissue and/or hemolymph sampled.

**Supplementary Fig. 14.**
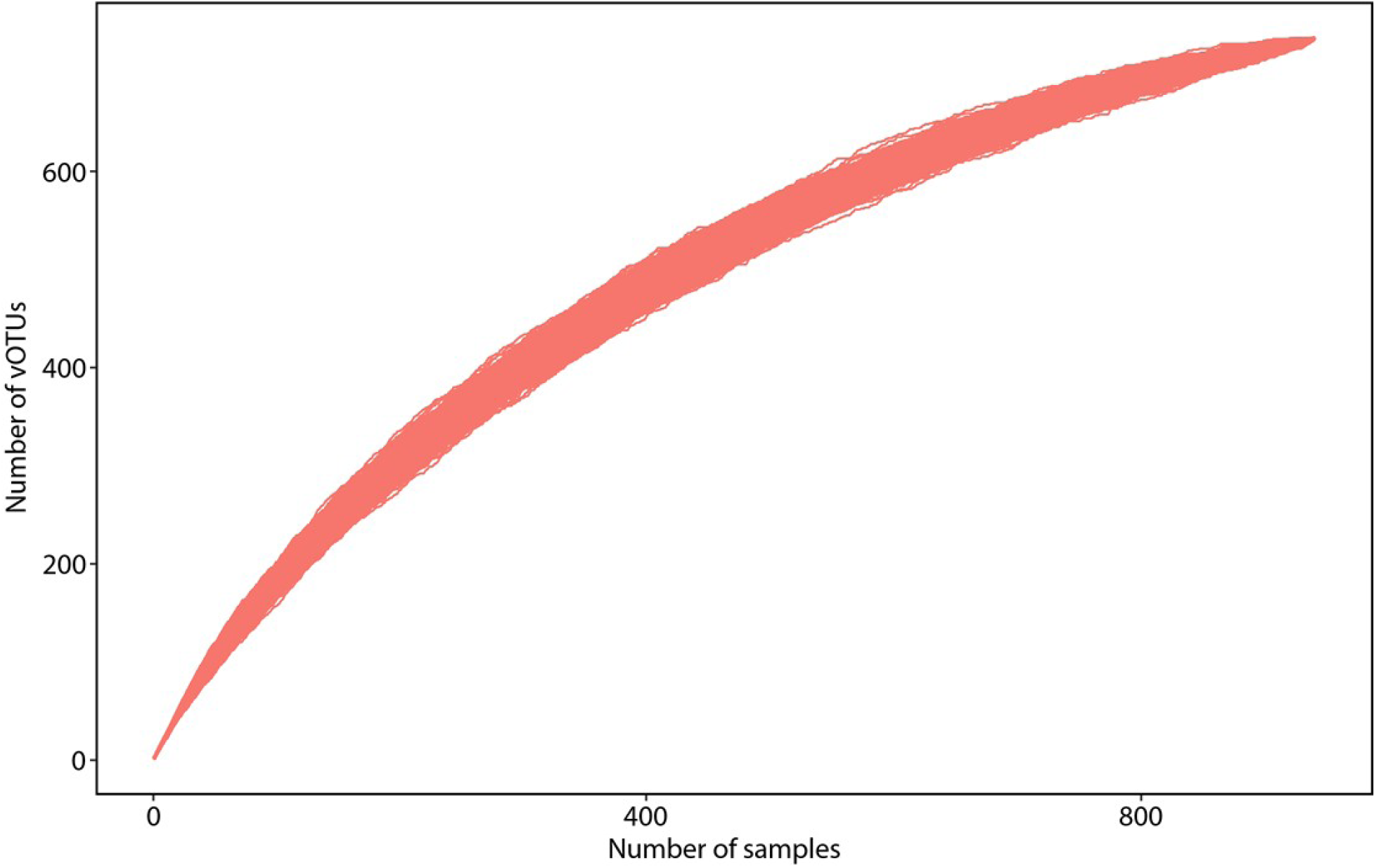
Rarefaction curve of RNA viruses detected in oysters. The rarefaction curves illustrate the predicted number of species-level viral Operational Taxonomic Units (vOTUs) detected in oysters (y-axis) as a function of the increasing number of transcriptomic samples (x-axis).

**Supplementary Fig. 15.**
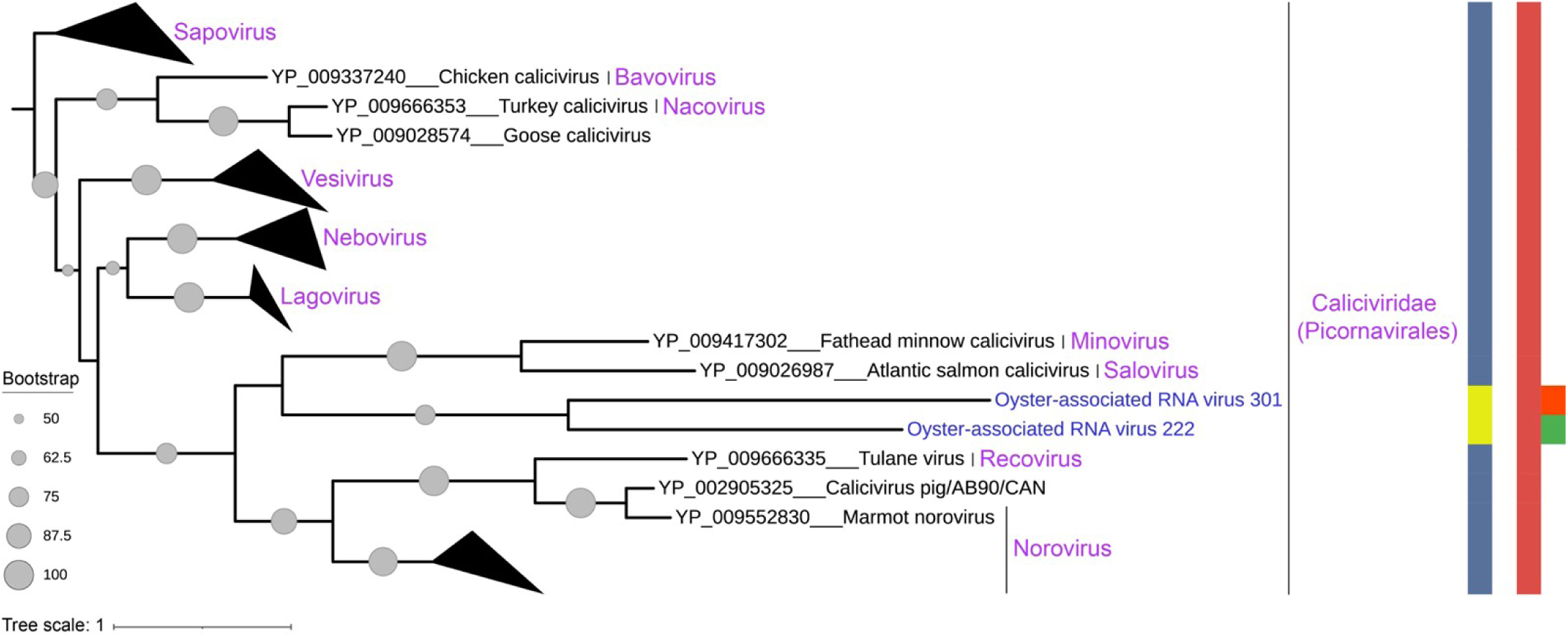
RdRp-palm phylogeny of oyster-associated RNA viruses (ORV) in the family *Caliciviridae* within the order *Picornavirales* from phylum *Pisuviricota*. The ML phylogenetic tree is constructed using RdRp palm sequences and re-rooted using members of the genus *Sapovirus*. The legend for the figure, including color key and tree annotations, is the same as in Supplementary Fig. 4.

## Notes

### Competing Interest Statement

The authors have declared no competing interest.

